# Directional Cell-to-cell Transport in Plant Roots

**DOI:** 10.1101/2024.09.16.613234

**Authors:** Léa Jacquier, Celeste Fiorenza, Kevin Robe, Jian-Pu Han, Fabienne Cléard, Christelle Fuchs, Priya Ramakrishna, Sylvain Loubéry, Linnka Lefebvre-Legendre, Marie Barberon

## Abstract

Cell-to-cell communication is critical for multicellular organisms. In plants, plasmodesmata—cytoplasmic channels—enable molecular transport between adjacent cells. In roots, this transport is predicted to be essential in nutrient acquisition and delivery to the vasculature. We demonstrate that plasmodesmatal transport persists in differentiated roots, despite apoplastic barriers such as Casparian strips and suberin lamellae in the endodermis, suggesting plasmodesmata as the sole pathway for water and nutrient flow at this stage. We also reveal a developmental switch in plasmodesmata function resulting in an unidirectional transport in differentiated roots. A genetic screen identified mutations that disrupt this directionality, leading to bidirectional transport. These mutations correlate with larger plasmodesmatal apertures, linked to defects in pectin composition and cell wall organization. This discovery underscores the role of plasmodesmatal aperture regulation and pectin in controlling directional transport. Our findings provide insights into plasmodesmata function and their regulation in roots.

## Introduction

Multicellular organisms rely on cell-to-cell communication and resource exchange to coordinate growth, development, and responses across tissues and organs. In plants, plasmodesmata form cytoplasmic connecting channels that enable the efficient redistribution of water, nutrients, and other macromolecules (including sucrose, hormones, as well as RNAs and proteins) ^1,2^. Plasmodesmata consist of a plasma membrane (PM) connection between neighboring cells across the cell wall, forming a pore containing an appressed structure of endoplasmic reticulum (ER) known as the desmotubule (DT). The space between the PM and DT, referred to as the cytoplasmic sleeve, allows the movement of molecules from one cell to another. The size, structure, protein, and lipid composition of the cytoplasmic sleeve will determine the types and sizes of molecules that can transit between cells^3^. Furthermore, the open and closed states of plasmodesmata can be regulated by controlling the deposition of the polysaccharide callose (beta1,3-glucan) within the cell wall, leading to a constriction of the plasmodesmata’s neck^2^. This plasmodesmata-mediated movement, referred to as symplastic transport, is involved in a multitude of biological process including defense against or susceptibility to pathogens, phloem loading and unloading and organ patterning^4–7^.

In roots, the symplastic transport is particularly well studied in the context of phloem unloading, where it plays a key role in post-phloem cell-to-cell transport at the root tip. Indeed, in undifferentiated roots, small molecules can move freely through plasmodesmata from phloem tissues down to the root meristem and outer root layers. This has been repeatedly observed by tracing phloem-mobile probes of different molecular weights applied to source leaves, such as CFDA (Carboxyfluorescein diacetate, 557 Da), or a GFP (Green Fluorescent Protein, 27 kDa) expressed in phloem companion cells with the *SUC2* promoter (*SUCROSE-PROTON SYMPORTER 2*) ^8–13^. Additionally, the presence of free symplastic transport in the root tip, was further supported by assays with the photoconvertible fluorescent reporter DRONPA (28 kDa) or microinjected Lucifer Yellow (444 Da) moving from the outer epidermis to the cortex, or by tracking the uptake of the symplastic tracer CFDA from the outer epidermis until the inner part of the root^14–16^. This symplastic continuum can be impaired by the deposition of callose in the cell wall of dividing and elongating cells. This was well demonstrated in the *cals3m* mutant (*callose synthase 3*), where CALS3 is constitutively active, as well as in plants overexpressing this mutation in a cell-type-specific manner^10,17,18^. Environmental conditions can also interfere with this symplastic transport in a callose-dependent manner, as shown for example, in the presence of mannitol or aluminum, excess iron, or in phosphate starvation in presence of iron^19–22^. Importantly, a common feature among the mutants and conditions tested, leading to an ectopic callose deposition in the root meristem, is that they are accompanied by dramatic effects in terms of meristem organization, division, and cell elongation, resulting in a halt in root growth. These root growth phenotypes can be partially explained by the lack of resource allocation to the meristem and interference with the transport of other molecules between root cell layers. Notably, the patterning of root cell layers is known to rely on cell-to-cell communication of signaling molecules and transcription factors (TFs), such as SHR (SHORTROOT) and SCR (SCARECROW) in the ground meristem^7,17,18,23^. Moreover, in recent years, it has become increasingly evident that plasmodesmata are also involved in hormonal transport, recently supported for auxin, ABA (abscisic acid), and brassinosteroid precursors^15,24–26^. Together, these studies are shaping a framework for understanding plasmodesmata regulation and function in the ground meristem.

In differentiated roots, plasmodesmata are predicted to play a major role in nutrient acquisition and transport from the root periphery (epidermis and cortex) to the inner vasculature^27,28^. Differentiated roots exhibit distinctive structures crucial for effective nutrient acquisition. For instance, root hairs on the outer epidermis enhance the surface area for nutrient absorption, while fully formed xylem vessels in the central vasculature facilitate the efficient redistribution of water and nutrients to various plant organs. Moreover, the endodermis (corresponding to the innermost cortical cell layer surrounding the central vasculature) forms barriers such as the Casparian strips (state I of endodermal differentiation) and suberin lamellae (state II of endodermal differentiation). These two endodermal barriers block respectively the transport through the apoplast (extracellular space) and the diffusion through the PM (transcellular) ^29–31^. Consequently, although undifferentiated roots theoretically allow free movement in the apoplastic space as well as in the symplasm along root cell layers, the presence of Casparian strips in the differentiated root endodermis compels transport exclusively through the symplasm. This concept was initially proposed by the work of de Rufz de Lavison in 1910, who investigated the penetration of various salts in living roots. He observed that salts unable to enter the root symplasm were obstructed in the apoplast at the level of Casparian strips. Meanwhile, salts entering the root symplasm were freely redistributed along the root tissues^32^. Nonetheless, evidence for a plasmodesmata-mediated transport in differentiated roots remain scarce. Its occurrence is mainly supported by observations with TEM (transmission electron microscopy) of plasmodesmata in the cell wall of differentiated root cells, including state I and II endodermis, in several species^33–38^. However, the presence of plasmodesmata does not necessarily guarantee a free symplastic transport of solute and macromolecules between differentiated root layers as described in the root tip. This can be illustrated with the case of root hairs, where symplastic tracer assays were performed. Indeed, in fully differentiated root hairs, where plasmodesmata are present, assays with DRONPA, or CFDA and Lucifer Yellow microinjections, indicated an absence of symplastic transport with their neighboring epidermal and cortical cells^14,16^. In this case, the mechanism restricting symplastic transport is not well understood but it is possible that smaller molecules such as water or nutrients would still be able to freely move from root hairs. Overall, we are still largely ignorant of whether symplastic transport can occur in differentiated roots and play a role in the radial transport from the outer epidermis to the inner vasculature.

Here we studied in detail the symplastic transport in undifferentiated *vs.* differentiated roots. We found that this transport occurs freely from the outer epidermal cells to the inner pericycle cells even in the presence of endodermal barriers. However, we observed that this transport becomes unidirectional and restricted from the cortex to the epidermis and from the endodermis to the cortex, as the root differentiates. This transport is impaired by osmotic stress suggesting that its driving force is the water flow. To decipher the mechanisms controlling this directional symplastic transport we performed a genetic screen aiming to identified mutants with a bidirectional symplastic transport in differentiated roots. We identified one mutant *ssm1 (sesame1*) with an enhanced and bidirectional transport. This mutant is impaired in rhamnogalactan synthesis in turn strongly affecting pectin and cell wall organization accompanied by defects in plasmodesmata. Altogether our data shed light on the previously understudied symplastic transport in differentiated roots, provides a better understanding of the role played by pectin in plasmodesmata and suggest a developmental transition in plasmodesmata directionality as the root matures.

## Results

### Unidirectional symplastic transport in differentiated roots

To study the symplastic transport along root differentiation and across root cell layers (Figure 1A), we first tested the capacity of roots to acquire and transport, from cell to cell, the symplastic tracer CFDA. We observed an uptake and redistribution of CFDA from epidermal to pericycle cells all along the root differentiation gradient including in the zones with Casparian strips (visualized with Propidium iodide block) ^29^ and patchy deposition of suberin (visualized with *GPAT5::SYP122-mScarletI*) ^31^ in the endodermis (Figure 1 B). To test if the symplastic transport from epidermal to pericycle cells could also be observed with a larger symplastic proxy we developed lines expressing the strongly enhanced GFP (sGFP2, monomeric, 26.9 kDa, maturation time 40 minutes) ^39^ in the root epidermis. To this end we used the previously described cell-type specific promoter *PDR9* (*PLEIOTROPIC DRUG RESISTANCE 9*) for the epidermis^40^ (Figure 1A,C and S1A). To be able to assess sGFP2 cell-to-cell transport in both undifferentiated and differentiated roots we used constitutive and estradiol-inducible promoters respectively. We observed that the epidermal-expressed sGFP2 accumulated in epidermal, cortical, endodermal and pericycle cells in both undifferentiated and differentiated roots (Figure 1D,E and S1B,C). This observation was corroborated by quantifications of sGFP2 fluorescence along root cell layers in both undifferentiated and differentiated parts of the root (Figure 1F). These results demonstrate that the symplastic transport can occur freely in differentiated roots and therefore comfort our current models of radial transport in roots.

**Figure 1:**
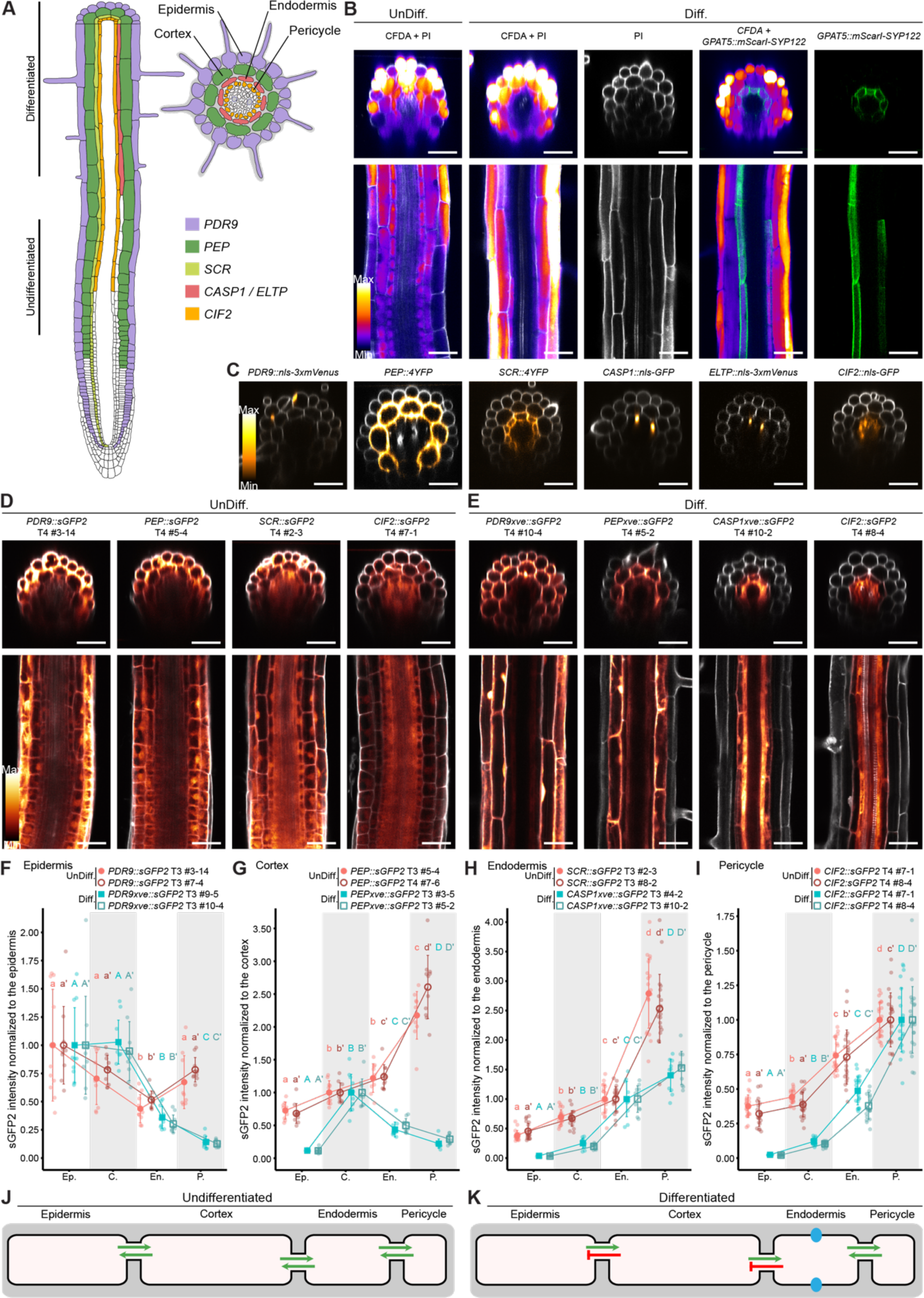
Directional symplastic transport in differentiated roots. (A) Schematic representation of root cell types and activity of cell type-specific promoters. Longitudinal and transversal views with the area of *PDR9*, *PEP*, *SCR*, *CASP1*, *ELTP* and *CIF2* promoters’ activities. Stages of endodermal differentiation are indicated, the undifferentiated zone corresponds to the elongating endodermis without endodermal barriers, while the differentiated zone corresponds to the zone with Casparian strips and patchy suberin deposition. (B) Live imaging of CFDA (4 hours staining) *vs.* PI uptake and CFDA uptake *vs. GPAT5::SYP122-mScarletI* signal. CFDA fluorescence is shown as a LUT (Look Up Table, Fire), and mScarletI signal is shown in green. (B, D, E) Undifferentiated (UnDiff.) and differentiated (Diff.) zones are shown, with transversal (upper panels) and longitudinal (bottom panels) views. (B-E) Cell walls counterstained with PI shown in gray, scale bars, 40 μm. (C) Live imaging of promoter-reporter lines in differentiated roots with transversal views. Reporter signals are shown as a LUT (Orange Hot). (D) Live imaging of sGFP2 expressed under cell-type specific promoters in undifferentiated zone (*PDR9* for the epidermis, *PEP* for the cortex, *SCR* for the endodermis and *CIF2* for the pericycle). (D-E) sGFP2 signal is shown as a LUT (Glow) for each construct, a second independent line is shown in Figure S1B,C. (E) Live imaging of sGFP2 expressed under inducible cell-type specific promoters in the differentiated zone (*PDR9xve* for the epidermis, *PEPxve* for the cortex and *CASP1xve* for the endodermis) and *CIF2* promoter for the pericycle. For inductions, seedlings were treated 19 hours with 10 μM β-estradiol. (F-I) Quantifications of sGFP2 relative intensity in the undifferentiated zone (circles) and the differentiated zone (squares), x-axis represents the cell layers analyzed (Ep. Epidermis, C. cortex, En. Endodermis, P. pericycle) and y-axis sGFP2 relative intensity. Each graphic displays two independents transgenic lines, for each zone, expressing the fluorophore in the epidermis (F), the cortex (G), the endodermis (H) or the pericycle (I). Quantifications are normalized to the fluorescence level in *UBQ10::3xsGFP2* (see Figure S1D, E) and presented relatively to the cell layer of production. Shapes and error bars represent the means and SD respectively, with overlaid dot plots representing each individual data (n≥10). Statistically significant differences (*P* < 0.05) between cell layers for each individual lines were tested using nonparametric Tukey test and are indicated by different letters. (J-K) Model illustrating symplastic transport in the undifferentiated (J) and differentiated zones (K). Green arrows represent free transport between cells, red marks the restriction of transport between cells and blue dots the Casparian strips.

Next, we wondered if this symplastic transport can also occur freely between the different root cell layers. To this end we used the same strategy and expressed sGFP2 constitutively or inductively in a cell-type specific manner with the previously described promoters: *PEP (PLASTID ENDOPEPTIDASE*) for the cortex^41^, *SCR* (*SCARECROW*) for the undifferentiated endodermis^41^, *CASP1* (*CASPARIAN STRIP MEMBRANE DOMAIN PROTEIN1*) and *ELTP* (*ENDODERMAL LIPID TRANSFER PROTEIN)* for the differentiated endodermis^31,42^ and *CIF2* (*CASPARIAN STRIP INTEGRITY FACTOR2*) for the pericycle and other stele tissues^43^ (Figure 1A,C and S1A). In undifferentiated roots, sGFP2 expressed in the cortex, the endodermis or the pericycle accumulated in all cell layers (Figure 1D, G-I, S1B) as expected for a free and bidirectional symplastic transport between root layers. However, in differentiated roots, we observed that sGFP2 expressed in the cortex accumulated in cortical, endodermal and pericycle cells but was not detected in the epidermis (Figure 1E,G and S1C). Moreover, sGFP2 expressed in the endodermis accumulated in the endodermal and pericycle cells but was not detected in the cortical and epidermal cells (Figure 1E,H and S1C). For the endodermis, the same result was obtained with a second promoter (*ELTP*) and two additional fluorophores: mScarlet and mScarlet-I (26.4 kDa, with maturation times of 174 and 36 minutes respectively) ^44^ expressed in the endodermis (Figure S1F,G). Furthermore, expression of *CASP1::cals3m*, leading to ectopic callose deposition in the differentiated endodermis^45^, in lines expressing sGFP2 in the differentiated cortex or endodermis lead to sGFP2 restriction to cortical and endodermal cells respectively (Figure S1H and S2A,B). Finally, we observed that sGFP2 expressed in the pericycle and stele accumulated in pericycle cells as well as in endodermal cells but was not detected in the cortical or epidermal cells (Figure 1E,I and S1C). Altogether these results demonstrate that in undifferentiated roots, cell layers are symplastically connected through a bidirectional symplastic transport while in differentiated roots, the symplastic transport becomes directional, occurring freely from the outer epidermis to the inner pericycle but is impaired from inner to outer cell layers at the epidermis-cortex and cortex-endodermis interfaces (Figure 1J,K).

### Water flow as a driving force for the symplastic transport in differentiated roots

Having observed that the symplastic transport occurs in differentiated roots in a directional fashion we wondered if this directionality could be explained by differences in callose deposition along the root cell layers or by the density or shape of plasmodesmata. Indeed, previous work focusing on directional symplastic transport suggested that all these factors could contributed to the unidirectional flow through plasmodesmata^10,14,46–48^. We first looked at callose deposition along the root differentiation gradient with Aniline blue staining as well by immunolocalization with an anti-callose antibody. In both case we could detect callose deposition at the cell plate in the meristematic region but could not detect any signal in the differentiated root of WT plants (Figure S2A,B). We next aimed to evaluate plasmodesmata density along root cell layers in differentiated roots. To this end, we intended to analyze the plasmodesmata marker PDLP5 (PLASMODESMATA LOCATED PROTEINS) in fully differentiated roots without the previously reported pleiotropic growth defects in lines expressing it under the *35S* promoter^26,49^. We therefore generated lines expressing PDLP5, fused to mScarletI expressed under the *UBQ10* promoter and successfully obtained plants without pleiotropic growth defects (Fig S2C). With these lines we observed a signal characteristic of plasmodesma in the meristematic zone, with punctuated signals at the cell periphery of very high density in the apico-basal side and of very low density in the outer-inner side (Fig S2D). In differentiated roots the signal was much weaker but following the same pattern and no difference was observed across root cell layers (Figure S2D). Finally, we analyzed plasmodesmata themselves at the ultrastructural level in differentiated roots with transmission electron microscopy (TEM). We were able to observe plasmodesmata between epidermal and cortical cells, between cortical and endodermal cells as well as between endodermal and pericycle cells even in the presence of suberin lamellae at the endodermis (Figure 2A). However, no structural difference was observed between cell layers, all plasmodesmata being similar to previous descriptions in undifferentiated roots^16,48,50^. Altogether these analyses did not allow us to pinpoint differences at the plasmodesmata level that could explain the directional symplastic transport observed in differentiated roots.

**Figure 2:**
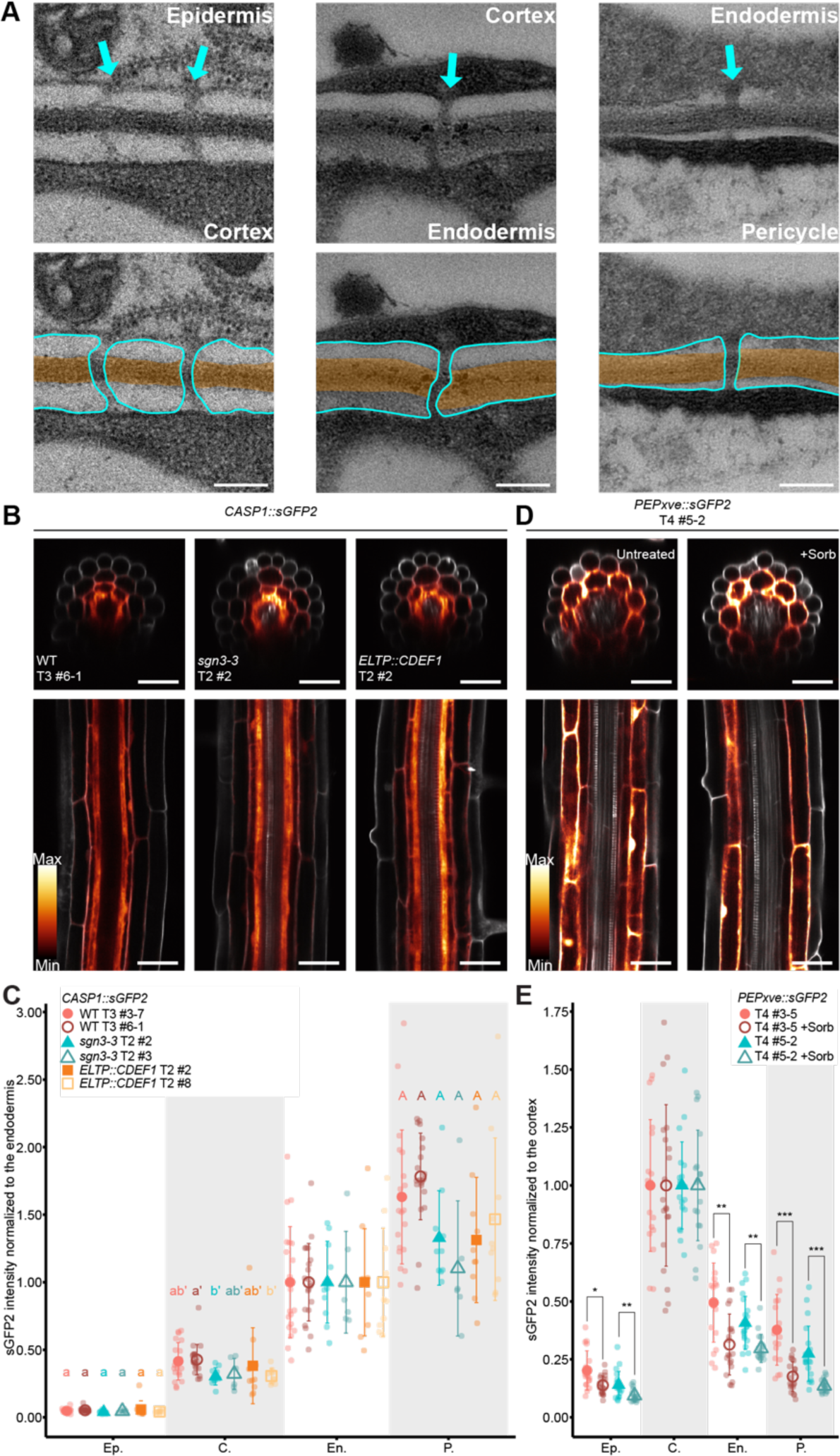
Water flow affects the symplastic transport in differentiated roots. (A) Transmission electron microscopy images of plasmodesmata across root cell layers in the differentiated zone, highlighted with blue arrows (cell-cell interfaces indicated in the upper images). Bottom panels represent the same pictures with overlayed drawing of the plasma membrane (in blue) and the cell wall (in yellow). Scale bars, 200 nm. (B) Live imaging of *CASP1::sGFP2* transgene in WT, *sgn3-3* and *ELTP::CDEF1* backgrounds in the differentiated zone. (B,D) sGFP2 signal is shown as a LUT (Glow), PI used to stain cell walls is shown in gray, transversal (upper panels) and longitudinal (bottom panels) views are shown, scale bars, 40 μm. For each construct, a second independent line is shown in Figure S2 E and F. (C) Quantifications of sGFP2 relative intensity of *CASP1::sGFP2* in WT (circles), *sgn3-3* (triangles) and *ELTP::CDEF1* (squares) backgrounds (n ≥ 6). (C, E) The x-axis represents the cell layers analyzed, and the y-axis, sGFP2 relative intensity. Quantifications are normalized to the fluorescence level in *UBQ10::3xsGFP2* (see Figure S1D, E) and presented relatively to the cell layer of production. Ep. epidermis, C. cortex, En. Endodermis, P. pericycle. Shapes and error bars represent the means and SD respectively, with overlaid dot plots representing each individual data. Statistically significant differences (*P* < 0.05) between lines at a given cell layers were tested using nonparametric Tukey test and are indicated by different letters. (D) Live imaging of sGFP2 signal in *PEPxve::sGFP2* line in untreated conditions (left panel) and after 2 hours of treatment with 150 mM sorbitol (right panel). For short induction, seedlings were treated with 10 μM of β-estradiol 2 hours prior to the sorbitol treatment. (E) Quantification of the relative sGFP2 intensity in the *PEPxve::sGFP2* line under untreated or sorbitol-treated condition as described in D (n ≥ 18). Differential statistical significance in each cell layer between untreated and sorbitol treatment are indicated for individual lines (\*\*\**, P* < 0.0005, ** *P* < 0.005, * *P* < 0.05) after nonparametric Tukey test.

In water transport models, endodermal barriers are predicted to play a central role in establishing a bulk flow for water allowing a directional transport from the root periphery to the vasculature as well as retention of water in the vasculature enabling a subsequent transport from roots to shoots^51,52^. These models are corroborated by analysis of water transport in Casparian strips and suberin deficient mutants and lines^53–55^. Moreover, we found that the directionality was established concomitantly with Casparian strips formation (Figure 1D-I). We therefore tested if functional endodermal barriers were required to establish the directional symplastic transport. To this end we analyzed a line expressing sGFP2 in the differentiated endodermis (with *CASP1* promoter) in the mutant *sgn3* (*schengen3/gassho1*, impaired in Casparian strips) ^56^ and in the suberin deficient *ELTP::CDEF1* line (*Endodermal Lipid Transferase::CUTICLE DESTRUCTING FACTOR1*) ^30,31^ but did not observe differences of sGFP2 accumulation across root cell layers in comparison to WT (Figure 2B,C and S2E).

Following the same rational, we next tested if interferences with water potential in the growth media would affect the symplastic transport in differentiated roots. To this end we added sorbitol in the growth conditions to create an osmotic gradient, thereby limiting water availability. Prior experiments showed that CFDA unloading in roots, was limited in presence of plasmolysing concentrations of sorbitol^57^. We therefore tested the effect of short sorbitol treatments, at a concentration not affecting root growth and cell viability, on sGFP2 movement between cells after 2 hours of estradiol-induction in *PEPxve::sGFP2* (Figure 2D,E and S2F, G). We observed that after 2 hours of 150 mM sorbitol treatment, sGFP2 remained confined in its cell type of expression (*ie.* the cortex), without inducing callose deposition (Figure S2H). Altogether our results comfort a model where the bulk flow of water would be the driving force for the symplastic transport in roots as previously suggested^15^.

### *ssm1,* a mutant with bidirectional and exacerbated symplastic transport

To identify cellular and molecular factors controlling this polarized symplastic transport in differentiated roots we performed a forward genetic screen. To this end, we EMS-mutagenized the line *ELTP::sGFP2* (parental line) where sGFP2 accumulates in endodermal and pericycle cells (see Figure S1F, G). After M1 propagation we screened in M2 for plants accumulating sGFP2 in outer root cell layers aiming to identify mutants with an inverted and/or bidirectional symplastic transport in differentiated roots. We called this screen and the identified mutants *sesame* (*ssm*) after the tale “Ali Baba and the forty thieves” and the phenotype of sGFP2 accumulating in outer layers, open sesame (Fig 3A). After screening with epifluorescent microscopy in M2 and confirmation with confocal in M3 we identified the open sesame mutant *ssm1-1*, where sGFP2 accumulated in all root cell layers (Figure 3B,C). We backcrossed *ssm1-1* with the parental line *ELTP::sGFP2* in M2, and observed in F2 that the open sesame phenotype was recessive, selected the corresponding seedlings and sequenced them. Sequence alignment to the parental line identified a region in chromosome 1 with mutations in 8 genes potentially causing the open sesame phenotype of *ssm1-1* (Table S1). Among these 8 genes, none of the corresponding loss of function mutants lead to an open sesame phenotype except *At1g78570* encoding for *ROL1* (*REPRESSOR OF LRX1*) also known as *RHM1* (*RHAMNOSE BIOSYNTHESIS 1*) (Figure S3A). The mutant *ssm1-1* harbors a missense mutation leading to a glycine-to-aspartic acid change at position 146 (Figure 3D). This *ssm1-1* mutant is allelic to the previously characterized *rhm1-4*, *rol1-1* and *rol1-2* mutants^58,59^ as well as to *ssm1-2* identified in our screen, which harbors a missense mutation leading to threonine 132 replaced by an isoleucine (Figure 3D, S3B,C). In addition to the open sesame phenotype, the mutant *ssm1-1* displayed a short root length, short root hairs and helicoidales flowers, phenotypes previously described in the allelic *rhm1* and *rol1* mutants (Figure S3D-G)^58–60^. Importantly, all these phenotypes including the open sesame phenotype were complemented by expressing *At1g78570* under its own promoter *(ROL1::ROL1g*) in *ssm1-1* (Figure 3B,C and S3D-H). Altogether our data identified *At1g78570* as the causal gene of the open sesame phenotype observed in *ssm1*. For consistency, we henceforth refer to the gene as *SSM1/ROL1* and to the mutants as *ssm1/rol1*.

**Figure 3:**
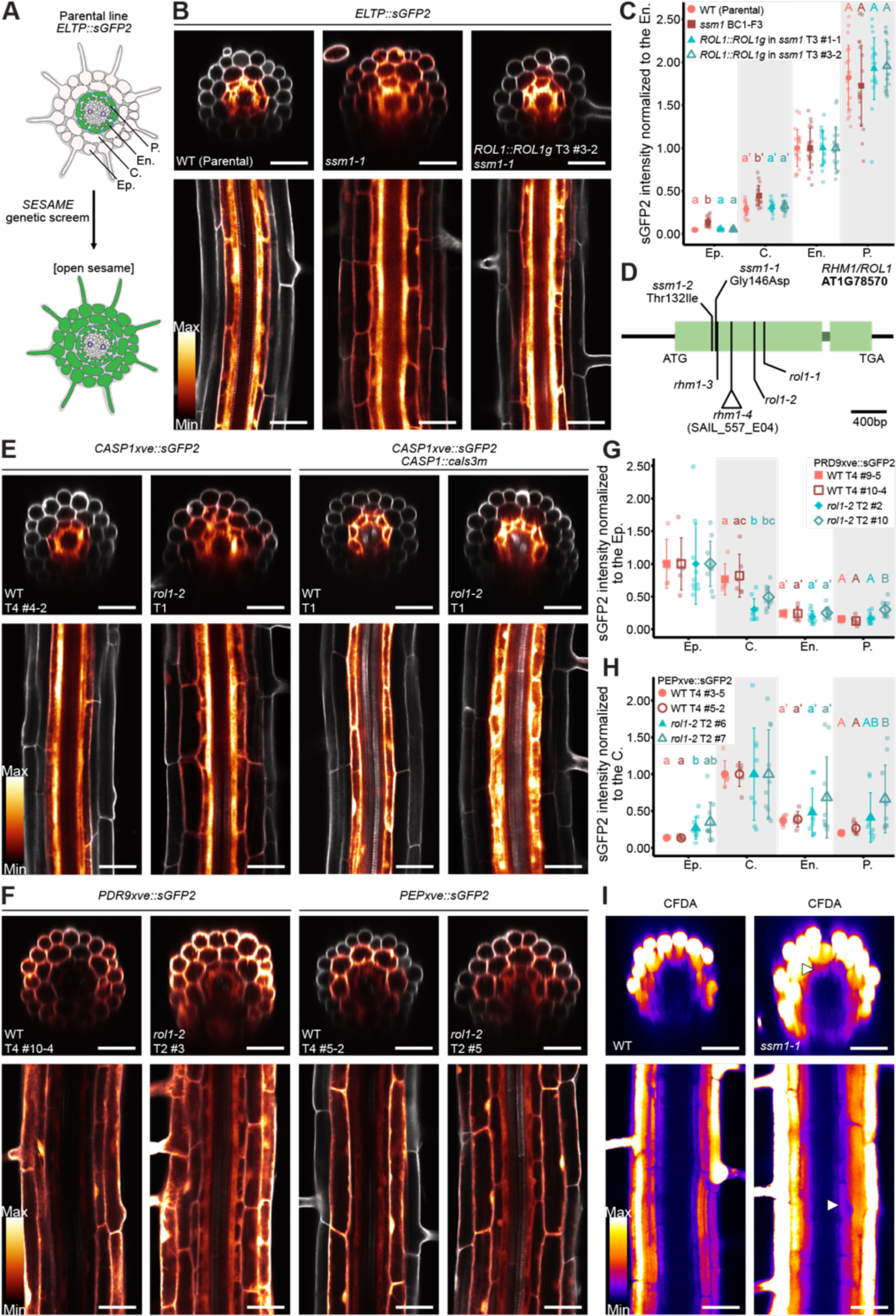
SESAME genetic screen identified a mutant with bidirectional and exacerbated symplastic transport. (A) Schematic representation of the *SESAME* genetic screen. Root cross-sections with sGFP2 signal, in green, representing the symplastic transport between the cell layers in the parental line (*ELTP::sGFP2* T4 #6-8) and the expected phenotype of mutants with bidirectional symplastic transport (*open sesame*). Ep. epidermis, C. cortex, En. endodermis, P. pericycle. (B) Live imaging of the *ELTP::sGFP2* transgene in WT, *ssm1-1* and *ROL1::ROL1g/ssm1-1* backgrounds in the differentiated zone. For the complementation with *ROL1::ROL1g,* a second independent line is shown in Figure S3H. (B, E, F, I) Transversal (upper panels) and longitudinal (bottom panels) views are shown. Scale bars, 40 μm. (B, E, F) sGFP2 signal is shown as a LUT (Glow); PI used to stain cell walls is shown in gray. (C) Quantifications of sGFP2 relative intensity from *ELTP::sGFP2* transgene in WT (circles), *ssm1-1* (squares) and *ROL1::ROL1g/ssm1-1* (triangles) backgrounds (n ≥20). (C, G, H) The x-axis represents the cell layers analyzed, and the y-axis shows sGFP2 relative intensity. Quantifications are normalized to the fluorescence level in *UBQ10::3xsGFP2* and presented relatively to average intensity in the cell layer of production. Shapes and error bars represent the means and SD, respectively, with overlaid dot plots representing each individual data. Statistically significant differences (*P* < 0.05) between genotypes at a given cell layers were tested using nonparametric Tukey test and are indicated by different letters. (D) Schematic representation of *At1g78570* locus with exons in light green and introns in dark green. Previously characterized alleles are indicated, as well as *ssm1-1* and *ssm1-2* identified in this study. (E) Live imaging of *CASP1xve::sGFP2* transgene in WT and *rol1-2* backgrounds with or without *CASP1::cals3m*, in the differentiated zone. For inductions, seedlings were treated 17 hours with 10 μM β-estradiol. (F) Live imaging of *PDR9xve::sGFP2* (left panels) and *PEPxve::sGFP2* (right panels) transgenes in WT and *rol1-2* backgrounds in the differentiated zone. For inductions, seedlings were treated 20 hours with 10 μM β-estradiol. (G-H) Quantifications of sGFP2 relative intensity of *PDR9xve::sGFP2* (G) and *PEPxve::sGFP2* (H) transgenes in WT and *rol1-2* backgrounds in the differentiated zone (n ≥5). (I) Live imaging of CFDA uptake (4 hours staining) in WT and *ssm1-1* roots in the differentiated zone. CFDA fluorescence is shown as a LUT (Fire) and arrows highlight the endodermal signal in *ssm1-1*.

We next wondered if the open sesame phenotype was the result of a bidirectional symplastic transport as initially intended by our screen. To this end, we first tested the expression of *ELTP::nls-3xmScarlet* in *ssm1-1* and observed an endodermis-specific signal indicating that endodermal identity was not affected in *ssm1-1* (Figure S3I). We next tested if callose deposition in the endodermis can suppress the open sesame phenotype as expected for a mutant with increased transport through plasmodesmata. To this end, we used the construct *CASP1::cals3m* leading to ectopic callose deposition in the differentiated endodermis^45^, combined with the inducible line *CASP1xve::sGFP2.* After estradiol induction, sGFP2 accumulated as expected, only in the endodermis and pericycle in WT roots and in all root cell layers in *rol1-2*, but was observed exclusively in the endodermis in both WT and in *rol1-2* mutant expressing *CASP1::cals3m* (Figure 3E). This demonstrated that *ssm1/rol1* is affected in the symplastic transport in differentiated roots. We next wondered if the symplastic transport was exacerbated in *ssm1/rol1* between other root layers. To this end, we expressed *PDR9xve::sGFP2* and *PEPxve::sGFP2* in *rol1-2* and observed, after estradiol induction, sGFP2 accumulating in all root cell layers (Figure 3F-H, S3J). Noteworthy, we could detect more sGFP2 signal in *rol1-2* compared to WT not only in the outer epidermis in *PEPxve::sGFP2* line but also more signal in the inner endodermis and pericycle with *PDR9xve::sGFP2* (Figure 3F-H, S3J). This suggest that the symplastic transport is not only bidirectional in *ssm1/rol1* but also exacerbated from the outer to inner root layers. To confirm this, we tested the uptake and distribution of CFDA in *ssm1-1* roots and observed an increased fluorescence in the endodermis compared to WT roots (Figure 3I). Altogether, these indicate that the symplastic transport is intensified in *ssm1/rol1* in both directions (*ie.* from the inner endodermis to the outer epidermis and from the outer epidermis to the inner pericycle).

### *ssm1,* a mutant with cell wall and plasmodesmata defects

*SSM1/ROL1* encodes a UDP-L-Rhamnose synthase involved in converting UDP-D-Glucose to UDP-L-Rhamnose. This activity was demonstrated *in vitro* ^61^ and was corroborated by cell wall analysis in the mutant displaying 40% reduction of rhamnose content^59^. Rhamnose plays a critical role in cell wall organization, particularly in the production of the pectin polysaccharides rhamnogalacturonan-I and rhamnogalacturonan-II (RG-I and RG-II respectively) ^62^. Previous work in the double mutant *lrx1-rol1* compared to *lrx1* (*leucine-rich repeat/extensin1*, a mutant whose root growth phenotypes are supressed by *rol1*) showed by immunolocalization on root sections in the root hair zone a strong reduction of labelling with the LM5 antibody (detecting RG-I) precisely in the cell wall of cortical, endodermal and vascular cells. This suggests a specific pectin defect in this mutant ,in specific cell types. In our experiments, using PI (Propidium Iodide) as a counter fluorescent staining for cell walls in roots, we consistently noticed that the signal was heterogenous in *ssm1/rol1* compared to WT, faint in the cell wall at the interface between two cells, and thicker and stronger at the cell corners (Figure 4A). PI binding pectin-containing cell walls, this observation was consistent with a broad pectin defect in *ssm1/rol1* in all root cell layers.

**Figure 4:**
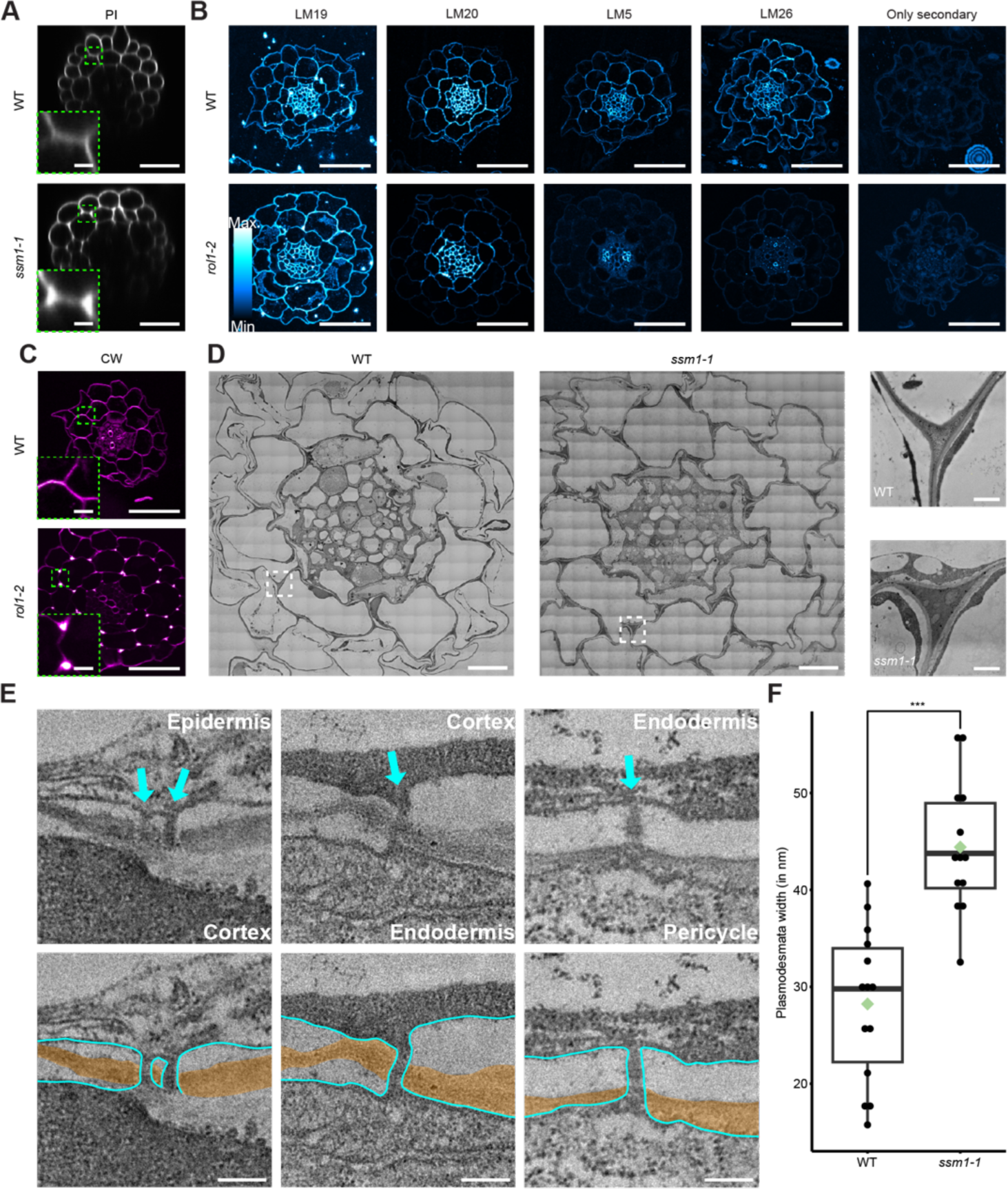
*ssm1,* a mutant with major cell wall and plasmodesmata defects. (A) Live imaging of PI staining in WT and *ssm1-1* (bottom panel) roots, in the differentiated zone, shown as cross section views. PI is shown in gray. Scale bars, 40 μm and 2 μm (closer views). (B) Cross sections of WT (upper panels) and *rol1-2* (bottom panels) after immunolocalization with LM19, LM20, LM5, LM26 antibodies and only the secondary antibodies, detecting respectively highly de-methylated pectin (HG), highly methyl-esterified pectin (HG), linear galactan (RG-I) and branched galactan (RG-I), shown as a LUT (Cyan Hot). Scale bars, 40 μm. (C) Calcofluor white staining in WT (upper panel) and *rol1-2* (bottom panel) from root cross sections in the differentiated zone. Calcofluor white signal is shown as a LUT (Magenta hot). Scale bars, 40 μm and 5 μm (closer views). (D) Transmission electron microscopy images of transversal root cross sections of WT (left panel) and *ssm1-1* (middle panel) in the differentiated zone and closer views of cell corners (right panels). Scale bars, 10 μm and 1 μm (closer views). (E) Transmission electron microscopy images of plasmodesmata across root cell layers in the differentiated zone in *ssm1-1* mutant. Plasmodesmata highlighted with blue arrows (cell-cell interfaces indicated in the upper images). Bottom panels represent the same pictures with overlayed drawing of the plasma membrane (in blue) and the cell wall (in yellow). Scale bars, 200 nm. See also Figure 2A for a WT comparison. (F) Quantifications of plasmodesmata width in WT and *ssm1-1*. Data presented as boxplots with overlaid dot plots, representing each individual data point and mean (green diamond), (n ≥ 13). Statistically significant differences were tested using a t-test and indicated (\*\*\**, P* < 0.0005).

Combined, our data suggested pectin defects in *ssm1/rol1,* beyond RG-I and in the wall of all root cell types. Consistently, analysis of *ROL1::nls-3xmVenus* lines showed that *SSM1/ROL1* promoter was active constitutively in all root cell layers as previously reported (Figure S4 A) ^58,63^. We therefore decided to investigate in more details the pectin defects in *ssm1/rol1* with immunolocalization on differentiated root sections with the antibodies LM19, LM20, LM5 and LM26 recognizing respectively unesterified homogalacturonan (HG), methyl-esterified HG, linear galactan side chains of RG-I, and branched galactan of RG-I^64–66^ (Figure 4B). LM19 labelled homogenously the cell wall across all cell types in WT but in *rol1-2* no signal was observed in the endodermal anticlinal walls and in the protoxylem and metaxylem walls (Figure 4B). LM20 antibody labelled the cell wall of stelar, cortical and endodermal cells (except the anticlinal wall of endodermal cells) in WT but in *rol1-2* the signal in the cell wall of cortical cells was strongly reduced (Figure 4B). LM5 labelled the cell wall across all cell types with a stronger signal in the phloem and a weak signal in the anticlinal endodermal walls and at the epidermis in WT roots, absent in *rol1-2* except at the phloem poles where the signal remained strong (Figure 4B). Finally, LM26 also labelled the cell wall across all cell types with a stronger signal in the phloem and no signal in the anticlinal endodermal walls in WT, absent in *rol1-2* with the exception of the metaphloem and cell corners in the stele (Figure 4B). Altogether, our data confirmed that RG-I are strongly reduced in *ssm1/rol1* in all root cell types as previously suspected but that HG are also affected especially in cortical and endodermal cells. Having observe defects in pectin beyond RG-I we wondered if the overall cell wall or cell wall modifications were affected in *ssm1/rol1*. We first could see with calcofluor white a strong defect in cellulose itself in *rol1-2* with a very faint signal in the cell wall between adjacent cells and a very strong ectopic signal in the cell corners similar to what we observed with PI (Figure 4C). Moreover, we could also detect ectopic callose deposition at the root periphery as well as at the cell corners of *ssm1-1* as described before for *rol1-2* (Figure S4B) ^67^. We next tested if other cell wall deposition in differentiated roots such as lignification and suberization were also affected in *ssm1/rol1.* Basic fuchsin staining for lignin did not unraveal differences between WT and *ssm1-1* in Casparian strips or in the xylem and fluorol yellow staining for suberin showed no defect in endodermal suberization (Figure S4C-E). Altogether our data indicate that *ssm1/rol1* harbors major but specific defect in its cell wall. Finally, we investigated *ssm1/rol1* at the ultrastructural level with TEM on root sections (Figure 4D). The anatomical organisation was comparable between WT and *ssm1-1.* However, the cell walls between adjacent cells were thinner and disorganized in the mutant (Figure 4D,E compared to Figure 2A). Moreover, the cell corners in *ssm1-1* were larger, denser and more heterogenous compared to WT between all root cell types (Figure 4D) a phenotype likely reflecting the pectin, and cellulose defects described above (Figure 4A, C). This ultrastructural phenotype was previously described in *rol1-2* and hypothesised to reflect a compensatory mechanism for cell wall defects in the mutant^67^. The mutant *ssm1/rol1* displaying symplastic defects, we wondered if plasmodesmata themselves would be affected in this mutant. To this end, we used TEM from root sections in the differentiated zone and checked all cell walls at the interface between cell types. Plasmodesmata were observed at the epidermis-cortex, cortex-endodermis and endodermis-pericycle interfaces and all displayed a structure comparable to WT plasmodesmata (Figure 4E and Figure 2A for WT comparison). However, plasmodesmata were on average ∼57% wider in *ssm1-1* compared to WT (Figure 4F). In summary, *ssm1/rol1* displayed major pectin and cell wall defects including cell wall and middle lamellae disorganisation together with an increase in plasmodesmata pore (Figure 5A). Such increase in plasmodesmata size could itself explain the exacerbated and bidirectional symplastic transport in *ssm1/rol1*.

**Figure 5:**
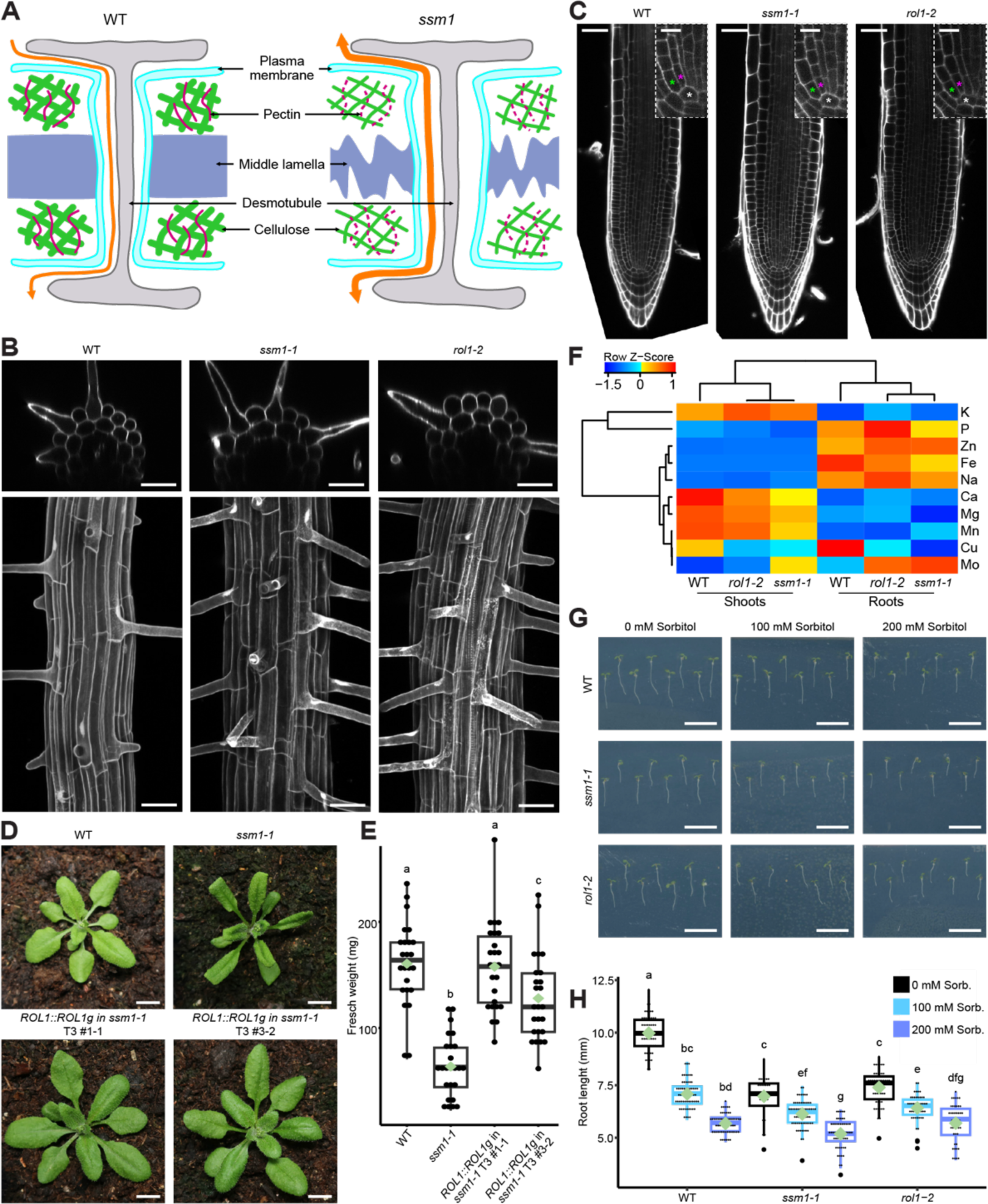
*ssm1*, a mutant with specific developmental and nutritional defects. (A) Schematic representation of a plasmodesmata in WT (left panel) and *ssm1/rol1* (right panel) with plasma membrane in blue, pectins in pink, middle lamellae in purple, desmotubule in gray, cellulose in green and cytoplasmic flow in orange. (B) 3D projection of PI staining in WT, *ssm1-1* and *rol1-2* seedlings to highlight the cell wall, transversal (upper panels) and longitudinal (bottom panels) views are shown. (B,C) PI is shown in gray; scale bars, 40 μm. (C) Root tip imaging in WT, *ssm1-1* and *rol1-2* seedlings stained with PI, and a close view of the meristematic zone, with white, green and magenta stars indicating the cortex-endodermis initial, the first cortical cell and the first endodermal cell respectively; scale bars, 10 μm. (D) Pictures of 3-week-old WT, *ssm1-1* and *ROL1::ROL1g* in *ssm1-1* grown in soil. Scale bars, 1 cm. (E) Quantifications of WT, *ssm1-1* and *ROL1::ROL1g* in *ssm1-1* fresh weight. Data presented as boxplots with dot plots overlaid, representing each individual data point and mean (green diamond), (n ≥ 24). Statistically significant differences (*P* < 0.05) between lines were tested using Tukey test and are indicated by different letters. (F) Heatmap based clustering analysis of the ionomic profiles in WT, *rol1-2* and *ssm1-1* shoots and roots. (See also Figure S5 A,B and Table S2 for numerical values). (G) Pictures of 5-day-old WT, *ssm1-1* and *rol1-2* seedlings growing on plates without, or with 100 mM or 200 mM sorbitol. Scale bars, 10 mm. (H) Quantifications of 5-day-old root length in WT, *ssm1-1* and *rol1-2* seedlings growing on plates without (black), or with 100 mM (light blue) or 200 mM sorbitol (purple). Data presented as boxplots with dot plots overlaid, representing each individual data point and mean (green diamond), (n ≥ 41). Statistically significant differences (*P* < 0.05) between lines and growing conditions were tested using nonparametric Tukey test and are indicated by different letters.

### *ssm1,* a mutant with specific developmental and nutritional

Having identified *ssm1*, a mutant with an exacerbated and bidirectional symplastic transport we wondered if this mutant could contribute to our understanding of plasmodesmata function in roots. As mentioned above and previously described, when grown on plates *ssm1* displayed a short root phenotype and short root hairs (Figure S3 F). We wondered if these phenotypes could be associated with defects in root patterning, a process highly regulated by cell-to-cell communication through plasmodesmata^7^. First, we looked in details at root hairs patterning in epidermal cells but could see that in the allelic mutants *ssm1-1* and *rol1-2*, root hairs formed in epidermal cells in contact with two cortical cells, resulting in an alternance of trichoblastic and atrichoblastic cell files at the root periphery comparable to WT plants (Figure 5B). Second, we carefully analyzed the ground meristem with a special focus on cortical-endodermal initial cells. We could observe some degree of disorganization in the division planes in *ssm1-1* and *rol1-2* however, the patterning in individual cell-files for the epidermis, endodermis and cortex was not affected (Figure 5C). In consequence, we could not observe cell-patterning defects in *ssm1*, its developmental phenotype likely being the consequence of reduced cell length (as can be seen in Figures 3B,E,F).

We next analyzed the mutant *ssm1* at the physiological level. After germination, *ssm1-1* plants were smaller but viable leading to fully developed adult plants, of reduced size but fertile (Figure 5D,E and Figure S3D-G). We used ionomics (where the mineral content of several elements is concomitantly analyzed in plant samples by ICP-OES, Inductively Coupled Plasma-Optical Emission Spectrometry) to evaluate the consequence of *ssm1/rol1* loss of function on mineral accumulation and translocation from root to shoots. To our surprise only minor differences could be observed between the allelic *ssm1-1 rol1-2* mutants and WT plants (Figure 5F and Figure S5 A-C). Nevertheless, the mutants had a similar and distinct ionome compared to WT plants, leading to significantly higher zinc and lower calcium levels in shoots and, lower iron level in roots. In turn, root-to-shoot translocation was reduced for calcium and increased for copper (Figure S5C). Finally, we tested the response of *ssm1*/*rol1* to water limiting conditions. First, we analyzed *ssm1-1* and *rol1-2* growth on plates in presence of sorbitol, and observed that contrarily to WT plants where the root length was strongly reduced (29 to 43% in 100 and 200 mM sorbitol respectively), the two allelic mutants were only slightly affected (12 to 26% in 100 and 200 mM sorbitol respectively) (Figure 5G,H). Second, we analyzed *ssm1/rol1* growth in response to drought in soil conditions. We observed that both *ssm1*/rol1 alleles had a higher recovery rate than WT plants after a long drought exposure followed by rewatering (Figure S5D,E). Altogether these results suggest a higher tolerance to drought stress in the mutant *ssm1/rol1,* with enhanced and exacerbated symplastic transport.

## Discussion

Our results reveal four significant findings. First, we confirmed predicted models of root transport by providing evidence that cell-to-cell transport through the symplast occurs along the root even after endodermal differentiation, when Casparian strips and suberin lamellae are present. Indeed, current models suggest that the symplastic pathway would be the only route available in the presence of apoplastic and diffusion barriers^27,28^. This prediction is supported by the presence of plasmodesmata in fully differentiated endodermal cells across various species^33–38^. However, to our knowledge, the occurrence of this pathway had not been previously demonstrated, as prior analyses of symplastic transport in differentiated roots focused on root hairs, where no transport was observed^14,16^. In this study, we observed plasmodesmata in fully differentiated endodermal cells of the model plant Arabidopsis and detected cell-to-cell movement from the outer epidermis to the inner pericycle, even in the presence of endodermal barriers. This observation was corroborated by both CFDA and GFP assays, which are classical proxies for cell-to-cell transport with molecular weight of 557 Da and 27 kDa, respectively. For context, among the 18 essential minerals^27^, the largest (molybdenum) has a molecular weight of 96 Da, metal-binding ligands (such as mugineic acid, nicotianamine, citrate, malate, histidine or phytate) ^68^ range from 134 to 660 Da, and the essential water-soluble vitamins (B1, B2, B3, B5, B6, B7, B9, and C) ^69^ range from 123 to 441 Da. Our results therefore indicate that the symplastic pathway remains freely open for passive diffusion of macro- and microelements essential for plant growth and development. This important validation will pave the way for future research to characterize the function of this pathway in nutrient acquisition and, more broadly, in cell-to-cell communication in differentiated roots.

Second, we found that while symplastic transport occurs freely from the outer epidermis to the inner pericycle, this bidirectional transport switches to a unidirectional mode as the root differentiates. Specifically, a free GFP expressed in cortical or endodermal cells was not detected in the outer epidermis and cortex, respectively. At a first glance, this result may seem surprising, given that plasmodesmata are often considered open gates for free diffusion between cells. However, from a physiological perspective, a system that allows nutrients - already selectively acquired at the root periphery through carriers - to be passively but efficiently transported in a directional manner without backflow would be particularly effective. Despite our efforts, we could not correlate this directionality with morphological changes at the plasmodesmata level. However, our result suggest that water flow is the driving force behind this symplastic transport, as previously suggested^51,52^. How this water flow becomes directional in differentiated roots remains to be clarified. Current models predict that bulk flow is established in the presence of the apoplastic barrier formed by Casparian strips. However, in our study we did not observe defects in directional cell-to-cell GFP transport in the absence of functional Casparian strips. Another possibility is that fully developed xylem vessels at the same developmental stage, along with root pressure and shoot transpiration create a suction effect. One limitation of our study is that the observed directionality is based on genetically encoded fluorophores (sGFP2, mScarlet and mScarlet-I) of the same size. Therefore, we do not yet know whether smaller molecules, including nutrients, are also transported directionally. Unfortunately, despite multiple attempts, we were unable to analyze cell-to-cell transport from inner to outer layers using a smaller proxy like CFDA in fully differentiated roots. In differentiated root cells, the large vacuole occupying most of the cytosolic space makes approach such as FRAP (Fluorescent Recovery after photobleaching) at the whole-cell level or microinjections extremely difficult without damaging or killing cells - two conditions that also induce callose deposition and would therefore compromise results. Nevertheless, our work demonstrates that molecules bellow 27kDa, moving passively through plasmodesmata, can travel from cell to cell but in a directional manner. This directionality likely has a significant implication not only for the efficient acquisition of nutrients and macromolecules for plant nutrition but also for signaling, where communication between the outer root layers and the root vasculature plays a crucial role. For example, this raises questions about how shoot-born signals controlling root responses are communicated, especially in the context of systemic nutrient signaling, where uptake capacity at the differentiated root periphery is highly regulated in response to shoot-derived signals (as well documented in systemic nitrate signaling^70^). In this context cell-to-cell communication between the vasculature and the outer epidermis might involve active transport of signaling molecules that overcome the directionality or perhaps a localized or transient loss of this directionality under specific conditions.

Third, we provide evidence that changes in plasmodesmata aperture can override this directionality. Through a genetic screen, we identified a mutant with bidirectional and exacerbated transport between root layers, accompanied by plasmodesmata with larger apertures. Notably, unlike the recently characterized *mctp3/4* and *mctp3/4/6* (*multiple c2 domains and transmembrane domain proteins*) mutants^50,71^, this effect can be reversed by callose deposition. Intriguingly, the *ssm1/rol1* mutant, which exhibits several growth defects related to changes in cell wall composition or flavonol rhamnosylation as previously described^58–60,63,67,72^, did not show major defects in biological functions involving cell-to-cell transport. This might reflect the ability of callose to fine-tune plasmodesmata-mediated transport in *ssm1/rol1* in specific cell types or conditions. Alternatively, it could indicate that plasmodesmata aperture is not the limiting factor for cell-to-cell communication and that other factors restrict this transport. An example of this is endodermal patterning, where plasmodesmata-mediated SHR movement from the vasculature is restricted to one cell layer due to its interaction with SCR^73^. Nevertheless, we observed moderate but specific alterations in mineral levels and reduced sensitivity to water-limiting conditions in *ssm1/rol1.* It is worth considering whether these phenotypes are associated with enhanced and bidirectional symplastic transport between epidermal and pericycle cells in *ssm1/rol1,* or if they are indirect consequences of the cell wall defects observed in this mutant. Our experiments do not allow us to differentiate between these two hypotheses. If symplastic transport itself is involved, we could envision a scenario in which exacerbated and bidirectional symplastic transport leads to largely futile transport between cell layers for most minerals, ultimately not affecting their translocation from roots to shoots, with xylem loading being the limiting step. An interesting case here is calcium whose cell-to-cell transport is thought to occur mainly though the endodermal symplast before xylem loading^74,75^. Recent evidence supporting this model includes the lack of effect of Casparian strips defects on shoot calcium content and the reduced calcium translocation to shoots in presence of ectopic suberin in the young endodermis^56,76^. In *ssm1/rol1,* we observed a reduction in calcium translocation from roots to shoots. This could indicate that, as previously suggested^56,76^, the limiting step for calcium transport is xylem loading, and that calcium needs to be concentrated in inner root layers for its efflux into the stelar apoplast. The increased growth observed in *ssm1/rol1* under water-limiting conditions or after rewatering could reflect an enhanced capacity to mobilize water in the mutant. Future work, particularly with additional mutants identified from the sesame screen, will help to test these hypotheses.

Finally, we identified a functional link between pectin and plasmodesmata. However, the mechanism by which the pectin and cell wall defects observed in the *ssm1/rol1* mutant affect plasmodesmata aperture remains unclear. One possibility is that the weakening and disorganization of the cell wall between adjacent cells in this mutant, along with compensatory ectopic deposition of pectin, cellulose and callose at the cell corners, alters the mechanical constraints of the cell wall surrounding plasmodesmata, leading to larger pores. Alternatively, the larger pores could indicate a more direct role of pectin, particularly RG-I and HG, in plasmodesmata. Supporting this hypothesis, several pectin-modifying enzymes were identified in plasmodesmatal proteomes^77,78^ and callose interactome^79^. Additionally, immunolocalization studies in tomato pericarp, apple fruits, and tobacco leaves showed that the cell wall around pit fields – area with a high density of plasmodesmata - is strongly enriched in specific pectin forms, especially unesterified HG^80–83^. More recently, enzymatic fingerprinting revealed specific pectin signatures in the cell wall purified with plasmodesmata, showing a strong enrichment of RG-I and the presence of RG-II and specific HG forms^84^. In light of these findings, pectin are expected to play a central role in plasmodesmata function. Current hypotheses suggest that pectin may influence plasmodesmata function and regulation by affecting pore elasticity and its callose-dependent regulation, or by impacting cell-cell adhesion, which in turn would affect plasmodesmata formation^84,85^.Since plasmodesmata create weak points in the otherwise solid cell wall, the larger pores in *ssm1/rol1* may reflect a significant role of pectin-mediated cell wall loosening in plasmodesmata formation.

Altogether, our data provide a framework to decipher the function of symplastic transport in roots, particularly in differentiated roots, where it has been largely understudied despite its expected major role. Our findings indicate that this transport occurs even in the presence of endodermal barriers, as predicted. Critically, it also demonstrates developmental transition of plasmodesmata transport as the root tissues differentiate from a bidirectional to a unidirectional symplastic transport. The sesame screen successfully identified a mutant impaired in this developmental transition and provided evidence for the role of pectin in plasmodesmata function. Future work with other sesame mutants will be equally valuable and is likely to identify new factors controlling plasmodesmata formation, regulation, and directionality. This will be crucial for gaining a better understanding of the mechanisms governing cell-to-cell transport and communication in developing and maturing plants.

### Experimental procedures

#### Plant material and growth conditions

All experiments were performed in Arabidopsis with the ecotype Col-0. Previously described T-DNA insertion lines and EMS mutants are the following: *rhm1-4^5S^* (SAIL_557_E04) , *rol1-1*, *rol1-2*^58^ and *sgn3-3*^56^. Previously characterized transgenic lines: *PEP::4YFP*, *SCR::4YFP*, *ELTP::nls-3xmVenus*, *CASP1::nls-GFP*, *CIF2::nls-GFP and CASP1::cals3m*^41–43,45^. Mutants and lines generated in this study are described below. For sterile plant culture, seeds were surface sterilized using chlorine gas (by combining 50/100 ml bleach with 3/6 ml 37% HCl for 2 hours). For *in vitro* experiments, seeds were sown in squared petri dishes containing half-strength MS basal salt mixture with 0.8% (w/v) Plant Agar, pH 5.7 and were stratified for 2-3 days in the dark at 4°C. All experiments on plates were performed with 5-day-old seedling unless otherwise specified. All plants were grown vertically under continuous light (∼100 µE) at 22°C. For propagation, transformation and physiological experiments, plants were sown on soil, stratified for 2-4 days at 4°C in the dark and then placed to grow in growth chambers under long day conditions (16 hours light, 8 hours dark) with LED lamps (120-150 µmol/m2/sec), at 20°C (+/- 2°C) and 55-75% relative humidity.

#### Plasmid generation and plant transformation

All Plasmids were generated using Multisite Gateway system. For non-destructive selection of transgenes, we used the destination plasmids pFR7m34GW (FAST red) and pFG7m34GW (FAST green) allowing fluorescence-based selection in seeds^86,87^. The following entry clones were previously described: *L4-pPRD9-R1*, *L4-pPEP-R1*, *L4-pSCR-R1*, *L4-pCIF2-R1*, *L4-pUBQ10-R1*, *L4-pCASP1-R1*, *L4-pELTP-R1*, *L4-GPAT5-R1* for promoters^30,40–43,45,88^, *L1-nls-3xmVenus-L2*, *L1-sGFP2-L2*, *L1-CDEF1-L2* and *L1-nls-3xmScarlet-L2* for coding sequences^30,39,45,89^, *R2-SYP122-L3*, *R2-mScarletI-L3* for C-terminal fusions^45,90^ and *pEN-R2-tHSP18.2-L3* and *pEN-R2-tNOS-L3* for terminators^86,91^. New entry clones were generated by Gibson assembly or BP recombination after PCR amplification into pDONR_P4–P1r or pDONR_P1-P2 or pEN-L4-ML-XVE-R1 (inducible system) ^92^. Details of the primers used for plasmid generation can be found in Table S3. The following clones were generated by Gibson assembly: *L4-pPDR9xve-R1* (*PDR9* promoter region 1949 bp upstream of the start codon, XhoI), *L4-pPEPxve-R1* (*PEP* promoter region 1490 bp upstream of the start codon, XhoI), *L4-pCASP1xve-R1* (*CASP1* promoter region 1207 bp upstream of the start codon, XhoI), *L4-pROL1-R1* (*ROL1* promoter region 1500 bp upstream of the start codon, BamHI), *L1-gPDLP1-L2* (*PDLP1* genomic region, NotI), *L1-gPDLP5-L2* (*PDLP5* genomic region, NotI), *L1-gROL1-L2* (ROL1 genomic region (2102 bp) and 600 bp of the downstream region, BglII/NotI). *L1-3xsGFP2-L2* was generated by amplification of sGFP2 and removing stop codons followed by a Gibson assembly. The following entry clones were generated through BP reaction: *L1-mScarlet-L2* and *L1-mScarletI-L2*.

All plasmids were generated following standard molecular techniques and confirmed through restriction digest and sequencing. DNA assemblies were performed using the NEBuilder enzyme mix (New England Biolabs E2621). Expression clones were introduced into electrocompetent Agrobacterium tumefaciens cells (strain GV3101) and subsequently transformed into Arabidopsis using the floral dip method^93^. For each transformation event, at least 10 T1 individuals were screened for expression and/or phenotypic traits. Experiments were performed on mono-insertional and homozygous T3 lines unless otherwise specified.

#### Physiological assays

For root imaging and root length quantifications, 5-day-old seedlings grown *in vitro* were imaged using Leica DM6 B microscope under polarized mode or scanned directly on plates with an Epson Perfection V800 flatbed scanner at 300dpi. Root length was determined by retracing roots in images from the root/hypocotyl junction to the tip using a Wacom Intuos Pro tablet and measured using Fiji^94^. Flower pictures of plants growing on soil were imaged using ZEISS Axio Zoom.V16 stereomicroscope. Whole plant pictures were imaged using Canon G7X camera. For sorbitol physiological assay, seeds were sown on ½ MS plates containing 0, 100 or 200 mM sorbitol, stratified for 2 days and place vertically to grow for 5 days, imaged and processed similarly as for root length measurement experiments. For drought treatment, water was withdrawn from 2-week-old plants growing in pots. The plants were grown for another 2 weeks without water. Pictures were taken at that time and 4 days after re-watering.

#### CFDA assay

CFDA was used as a symplastic tracer as described before with minor modification^15^. Seedlings were placed for 4 hours in 5 µg/ml CFDA diluted in water in the dark and rinsed in water 10 seconds prior to imaging.

#### Cell wall staining

Propidium iodine (PI) was used to visualize the apoplastic space and to visualize cell walls. Seedlings were live stained in 15 µM PI, diluted in water, and kept in the dark for 5 minutes prior to imaging. Callose was stained using aniline blue. Seedlings were fixed with 4% formaldehyde dissolved in 1X phosphate-buffered saline (PBS) for 30-40 minutes. After removing formaldehyde, seedlings were rinsed 3 times with 1XPBS for 10 minutes. At this stage, seedlings were stored in 1XPBS for up to 2 weeks. Seedlings were then stained overnight with a 0.01 mg/ml aniline blue solution (Biosupplies, Cat# 100-1) and washed with 1XPBS 30 minutes before imaging. Whole seedlings were mounted on glass slides with water. Suberin was stained using fluorol yellow. Seedlings were placed in a solution of FY088 (0.01% wt/vol, lactic acid) (Chem Cruz, Cat# sc-215052) for 30 minutes at 70°C and washed twice with water. Seedlings where counterstained with Aniline blue (0.5% wt/vol, water) (Sigma-Aldrich, Cat# 415049) for 30 min and washed 3 times with water. Whole seedlings were mounted on glass slides with 50% glycerol and observed using an epifluorescence stereomicroscope Leica M205 FCA with a GFP filter excitation: 450 to 490 nm, emission: 500 to 550 nm. Quantifications of the root length of three different suberin patterns (non-suberized, patchy and suberized) were done in Fiji as previously described^89^. Basic fuchsin staining was performed on cleared roots to visualize lignin as previously described^95^. Seedlings were fixed with 4% formaldehyde in 1XPBS for 1 hour. After 2 washed of 1 minute in 1XPBS, seedlings were transferred to a ClearSee solution containing 10% xylitol, 15% sodium deoxycholate and 25% Urea, overnight. After clearing, seedlings were stained with 0.2% Basic Fuchsin dissolved in ClearSee solution overnight. All incubations steps are performed in the dark, at RT and under gentle agitation. Prior to observation, seedlings were washed every 30 minutes for 5 hours with ClearSee solution. Whole seedlings were mounted on slides with ClearSee solution.

#### Immunolocalization

The following concentrations of antibodies were used: LM5, LM19, LM20 and LM26 1:10; (1→3)-β-Glucan (callose) 1:400; Alexa Fluor Plus 488 (Goat anti Rat) 1:500; Alexa Fluor Plus 488 (Goat anti Mouse) 1:200. For whole-mounted samples, seedlings were fixed with 4% formaldehyde supplemented with 0.1% triton, in a desiccation jar under vacuum for 2-3 minutes, twice, followed by 1 hour without vacuum. Samples were washed 3 times with 1XPBS for 10 minutes after which 4-5 seedlings were placed vertically on “superfrost” slides. After drying overnight, a counter-slide was added to the slides in 1XPBS. An automated *in situ* hybridization and immunohistochemistry system: InsituPRo (© CEM Corporation) was used for the following steps. Tissues were rehydrated 10 times with 1X-PBS with 0.05% Triton for 10 minutes. The cell wall was digested with 2% driselase in 1XPBS for 90 minutes at 37°C, followed by 3 washes with 1X-PBS with 0.05% Triton for 10 minutes. Then, the membranes were permeabilized with 1XPBS, 10% DMSO and 3% IGEPAL for 30 minutes at RT twice, followed by 4 washes with 1X-PBS with 0.05% Triton for 10 minutes. Samples were blocked for 1 hour with 1XPBS and 2% BSA. Incubation with primary antibodies was performed for 4 hours at 37°C in 1XPBS and 2% BSA followed by 5 washed with 1X-PBS with 0.05% Triton for 8 minutes. Incubation with secondary antibodies was performed for 3 hours at 37°C, followed by 5 washes with 1X-PBS with 0.05% Triton for 8 minutes. Cellulose was stained with 1% calcofluor white, to visualize the cell walls, for 5 minutes and washed 5 times for 5 minutes with 1XPBS. Samples were mounted using Mowiol and stored at 4°C until imaging at the confocal. For semi-thin plastic-embedded sections, seedlings were fixed in 4% formaldehyde for 1 hour at room temperature and washed with 1XPBS. Seedlings, aligned at the root tip, were placed by group of 4 to 5 in 2% agarose blocks. Dehydration steps were the following: 30% ethanol for 40 minutes, 50% ethanol for 40 minutes, 70% ethanol for 40 minutes, 100% ethanol for 1 hour, then overnight in 100% ethanol. For infiltration, agarose blocks were placed in 30% LR white (diluted with 100% ethanol) for 4 hours, 70% LR white for 4-8 hours and finally overnight in 100% LR white. Each block was placed in PTFE mold (TED PELLA INC, 10506), submerged with 100% LR white and sealed, then incubated 24-48 hours at 60°C for polymerization. Sections of 2 μm were made using HistoCore AUTOCUT Leica microtome and placed on “superfrost” slides. Immunolocalization was performed with InsituPRo robot. Samples were rehydrated twice with 1X-PBS with 0.05% Triton for 10 minutes, then blocked for 30 minutes with 2% BSA diluted in 1XPBS. Primary antibodies in 1XPBS and 2% BSA were incubated for 1 hour at 37°C followed by 5 washes of 5 minutes in 1XPBS. Secondary antibodies in 1XPBS and 2% BSA were incubated for 1 hours at 37°C followed by 5 washes of 5 minutes in 1XPBS. Cellulose was stained with 1% calcofluor white, to visualize the cell walls, for 5 minutes and washed 5 times for 5 minutes with 1XPBS. Samples were mounted using Mowiol and stored at 4°C until imaging at the confocal.

#### Pharmacological treatment

For inducible transgenic lines, 5-day-old seedlings growing on ½ MS agar plates were sprayed with 10 µM liquid β-estradiol, solubilized in DMSO, diluted 1/1000 in liquid ½MS. The duration of the treatment is specified for each experiment. For short term sorbitol experiments, seedlings were grown for 5 days on ½ MS agar plates, sprayed with 10µM liquid β-estradiol, solubilized in DMSO, diluted 1/1000 in liquid ½MS, for 2 hours and then transferred to ½ MS agar plates with 150 mM sorbitol, and imaged two hours after transfer.

#### Confocal microscopy

Most of the imaging were taken in the zone where cells are still in elongation phase referred as undifferentiated zone or in the zone where Casparian strips are present and where the endodermis is partially suberized (patchy zone) referred as differentiated zone. Confocal laser-scanning microscopy experiments were performed on a Leica SP8. Excitation and detection parameters are the following: sGFP2/GFP/YFP/mVenus 488nm, 498-536nm; mScarlet/mScarletI 552nm, 593-642nm; mCherry 552nm, 600-640nm; mCitrine 488nm, 520-560nm; Gamillus 488nm, 510-540nm; PI 488nm, 604-659nm; CFDA 488nm, 500-550nm; Aniline blue 405nm, 410-470nm; Basic fuchsin 552nm, 600-650nm; Alexa Fluor Plus 488 488nm, 520-548nm. Exept for immunolocalization and calcofluor white(see above), all the fluorescent images representing cross sections were obtained in live as virtual cross sections acquired in xzy with the microscope. Imaging of *UBQ10::PDLP5-mScarletI* line was performed on a Leica Stellaris 8 FALCON with the following excitation and detection parameters: 552nm, 596-642nm. The images were deconvoluted using integrated Leica lightning. Images were imported in Fiji for further analyses, as well as 3D projections if needed and for LUT (Look Up Table) application.

#### Fluorescence’s quantification and analysis

Images were analyzed with Fiji (https://fiji.sc/). Quantifications of the fluorescent intensity in each cell layers were performed on xzy images using a semi-automatized ImageJ macro consisting of checking the absence of saturation was checked using the “Hilo” LUT. After this, a region of interest was drawn in the background to subtract it from the biological signal. Next, in each cell layers, for the upper part of the root, a line was drawn allowing to calculate the average intensity in each cell layer. Data were normalized as follow. First, to minimize the influence of cell size and intracellular content, a coefficient was determined in *UBQ10::3xsGFP3* transgenic line, in the undifferentiated and the differentiated zone in each cell layers and applied to the data. This coefficient is presented in Figure S1 D-E. Second, to compensate for differences in promoter activity and potential variation in expression between independent lines, the average signal of the cell layers of production was set to 1.

#### Electron microscopy

Seedlings were fixed in 2.5% glutaraldehyde in 1XPBS with an addition of 0.1% tween for 1-2 hours at RT. Fixator was replaced by 2 ml of osmium in 1XPBS with 1.5% of potassium hexacyanoferrate (II) trihydrate for 1-2 hours. Seedlings were rinsed with 1XPBS until the solution appear clear. Seedlings, aligned at the root tip, were placed by group of 4 to 5 in 2% agarose. Blocks were dehydrated with 30% ethanol for 40 minutes, 50% ethanol for 40 minutes, 70% ethanol for 40 minutes, 100% ethanol for 1 hour and overnight in 100% ethanol. The infiltration was done with 30% LR white (diluted with 100% ethanol) for 4 hours, 70% LR white for 4-8 hours and 100% LR white overnight. Finally, the blocks were placed in PTFE mold (TED PELLA INC, 10506), submerged with 100% LR white and sealed, then incubated 24-48 hours at 60°C for polymerization. Blocks were cut using Leica UCT Ultracut Ultramicrotome. Ultrafine sections were made for 2D analysis (80-60 nm). The sections were placed on single slot formvar coated grids (25-50 nm Formvar and 3-4 nm Carbon, Electron Microscopy Sciences). Grids were stained for 10 minutes with 2% uranyl acetate, rinsed with water, incubated 3 minutes with lead citrate (Renolds) and rinsed with water. Images were acquired on a Talos L120C (120 KeV, LaB6) electron microscope. Image analysis was performed with IMOD^96^ and Fiji^94^.

#### Genetic screen

Approximately 25,000 seeds of *ELTP::sGFP2* T3 #6-8, parental line, were soaked overnight at 4°C in 40 ml of 100 mM PBS. Seeds were treated with 0.4% EMS diluted in 100 mM PBS for 8 hours under gentle mixing, then washed 20 times with 1XPBS. Seeds were resuspended in 0.15% agar solution for ease of sowing and placed in the dark at 4°C for 2 days. Seeds were sown on soil. M1 plants were harvested individually, yielding almost 3024 M2 populations. M2 seeds were surface-sterilized and sown on ½MS agar plates grown vertically. Plates were kept at 4°C in the dark for around two days before placing in growth chambers. 5 days-old independent M2 lines were screened, with 25 seeds for each. Screening was done directly on plates using a Nikon Eclipse 80i wide field microscope (10X PlanFluor objective NA 0.3. with GFP filter 457-487nm, 502,537nm). Individuals with a putative mutant phenotype (open sesame) were transferred to soil and propagated. The open sesame phenotype was confirmed in M3 on 5-day-old seedlings grown vertically with a Leica SP8 confocal microscope. In parallel, selected M2 plants were backcrossed with the parental line. Segregation counts were performed on the resulting F2 from which 20 seedlings with a clear open sesame phenotype were collected and pooled for subsequent DNA extraction. An expected mean coverage of approximately 50x for each genome was expected. Genomic DNA was extracted using the DNeasy Plant Mini Kit (Qiagen, 69104). Whole-genome sequencing libraries were prepared by the iGE3 (Institute of Genetics and Genomics of Geneva). Libraries were multiplex six times for single-end sequencing. All mutants were mapped against DNA library obtained from a pool of 20 seedlings from the parental line. Alignments of the small reads (100bp) to the Arabidopsis reference genome, as well as comparisons between mutant and parental pools, were performed using the SIMPLE fully automated pipeline (https://github.com/wacguy/Simple)^97^. Alongside, F2 seedlings were crossed with WT to obtain outcrossed plants containing *ssm1-1* mutation without the initial transgene (*ELTP::sGFP2*), homozygous plants were obtain in F2 and used for further analysis, they are referred to as *ssm1-1 outcrossed (ssm1-1^oc^).* This *ssm1-1^oc^* was used for the experiments presented figures 3I, 5B,C,F,G and for Figure S5 where *ssm1-1* is written for simplification. Experiments with *ssm1-1* were performed with the parental line as WT.

#### Ionomics

9 days old seedlings were used for the quantification of metal concentrations. Shoots and roots were harvested separately. Roots and shoots were desorbed by washing in 10 mm Na2EDTA for 10 min and rinse three time with ultrapure MiliQ water for 1 min. After desorbing, tissues were dried using paper towels and put at 65° degrees. After drying, between 5 to 10 mg of dry tissue per biological replicate were mixed with 750 ml of nitric acid (65% [v/v]) and 250 ml of hydrogen peroxide (30% [v/v]). After one night at room temperature, samples were mineralized at 85°C for 24 hours. Once mineralized, 4 ml of milliQ water was added to each sample. Metal contents present in the samples were then measured by Inductively Coupled Plasma Optical Emission Spectrometry (ICP-OES). PCA, Variable factor maps and Row Z-score data were generated using MVapp^98^. Shoot to root ratios were obtained by dividing the average concentration of each mineral in shoots by its average concentration in roots.

#### Graphical representations and statistical analysis

Quantitative graphics were generated using RStudio with the ggplot package. Data analysis was conducted within the R environment (R Core Team 2023). Dotplots represent the mean value (with various shapes), the standard deviation and dots for each individual data point. All boxplots represent the median value (line), upper and lower quartile (box), the mean value (diamond), the interquartile range (whiskers) and dots representing each individual data point. Statistical analysis was also performed in R, with sample sizes (n) specified for each experiment. Statistically significant differences are indicated by different letters for multi-comparison tests or asterisks for pair-wise comparisons. The legend for each figure panel containing quantifications provides brief descriptions of the statistical analyses and significance levels. Prior to comparisons, normality was tested using Shapiro test and homoscedasticity using Var.test, Bartlett test or Levene test depending on data characteristics. For data following parametric conditions, Student test was used for single comparison and anova follow by a parametric Tukey test for multiple comparison. For multiple comparisons of data not following parametric conditions, non-parametric Kruskal-Wallis and non-parametric Tukey tests were used. Graphical representations were compiled in R, figures and illustrations were compiled with Affinity.

## Acknowledgments

We thank Tonni G. Andersen, Satoshi Fujita, Joop Vermeer, Lothar Kalmbach and Niko Geldner for sharing plasmids for MSG. Joop Vermeer and Christoph Ringli are thanked for sharing seeds. We thank the Electron Microscopy Facility (EMF) of the University of Lausanne for help with TEM in particular Jean Daraspe for advice with TEM and Damien De Bellis for advice and for sharing protocols for TEM. We thank Mylene Docquier and members of the iGe3 (Institute of Genetics and Genomics in Geneva) at the University of Geneva for help with NGS and Christoph Bauer and Jérôme Bosset from the Photonic Biomaging Center as well as Yashar Sadian and Andrew Howe from DCI (Dubochet Center for Imaging) at the University of Geneva for help with confocal and electron microscopy respectively. We thank Sandrine Chay and Stephane Mari from the Multi-Elemental Analyses Service (SAME) at the Institute for Plant Sciences of Montpellier (IPSiM). We would like to thank Isabelle Fleury for help with the propagation of plants as well as the mutagenized population. Maria Beatriz Capitão is thanked for performing preliminary test with suberin staining. We thank Eliott Larue and Charlotte Grometto for their help during the design of the symplastic tracking under sorbitol treatment. Lothar Kalmbach is thanked for multiple advice and critical inputs along this project especially for the genetic screen and for critical reading of this manuscript. This work was supported by iGe3 (Institute of Genetics and Genomics in Geneva) with a PhD salary award to LJ and by funding from the Sandoz Family Monique De Meuron philanthropic foundation’s program for academic promotion and the Swiss National Science Foundation (Grant 31003A_179159) to MB and by the state of Geneva.

