## Supplemental Figure 1 for "Directional Cell-to-cell Transport in Plant Roots"

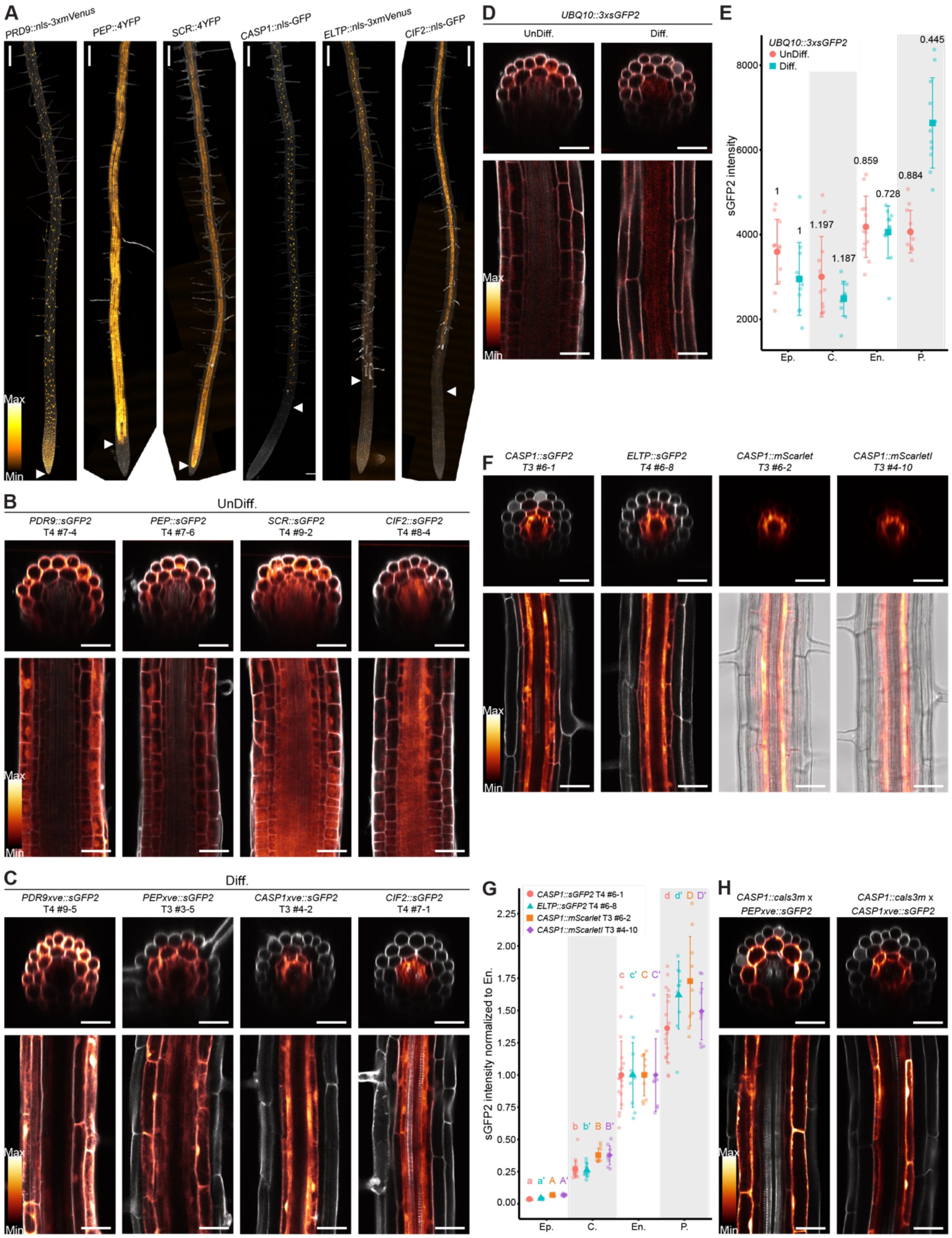


**Figure S1: Directional symplastic transport in differentiated roots.**

(A) Live imaging of promoter-reporter lines (tile images with z-projection), fluorophores shown as a LUT (Orange hot). White arrows highlight the signal onset. Scale bars, 200 μm. (A-D, F, H) Cell walls counterstained with PI shown in gray. (B) Live imaging of sGFP2 expressed under cell-type specific promoters in undifferentiated (UnDiff.) zone (*PDR9* for the epidermis, *PEP* for the cortex, *SCR* for the endodermis and *CIF2* for the pericycle) (Related to Figure 1D, F-I). (B-D, F, H) Transversal (upper panels) and longitudinal (bottom panels) views are shown, sGFP2 signal is shown as a LUT (Glow), scale bars, 40 μm. (C) Live imaging of sGFP2 expressed under inducible cell-type specific promoters in the differentiated (Diff.) zone (*PDR9xve* for the epidermis, *PEPxve* for the cortex, *CASP1xve* for the endodermis) and *CIF2* promoter for the pericycle (Related to Figure 1E, F-I). For inductions, seedlings were treated 19 hours with 10 μM β-estradiol. (D) Live imaging of *UBQ10::3xsGFP2* in the undifferentiated (left panel) and differentiated zone (right panel). (E) Quantifications of *sGFP2* signal in *UBQ10::3xsGFP2* in the undifferentiated (shown with circles) and in the differentiated zone (shown with squares). The x-axis represents the cell layers analyzed, and the y-axis sGFP2 signal intensity. Shapes and error bars represent the means and SD respectively (n ≥11). Numbers above corresponds to correction coefficient applied to all further quantifications relatively to the epidermis. (E,G) Ep. epidermis, C. cortex, En. Endodermis, P. pericycle. (F) Live imaging of *CASP1::sGFP2*, *ELTP::sGFP2*, *CASP1::mScarletI* and *CASP1::mScarlet,* in the differentiated zone. mScarlet and mScarletI signals are shown as a LUT (Glow) overlaid with bright field (right, bottom panels). (G) Quantifications of sGFP2, mScarlet and mScarletI signal intensities in *CASP1::sGFP2*, *ELTP::sGFP2*, *CASP1::mScarlet* and *CASP1::mScarletI* lines in the differentiated zone. The x-axis represents the cell layers analyzed, and the y-axis shows the fluorophore intensity. Quantifications are normalized to the fluorescence level in *UBQ10::3xsGFP2* and presented relatively to the cell layer of production. Shapes and error bars represent the mean values and SD respectively, with overlaid dot plots representing each individual data (n ≥ 20). Statistically significant differences ($P < 0.05$) between cell layers for each individual lines were tested using nonparametric Tukey test and are indicated by different letters. (H) Live imaging of sGFP2 in F1 from crosses between lines expressing *CASP1::cals3m* and sGFP2 under inducible cell-type specific promoters (*PEPxve* for the cortex and *CASP1xve* for the endodermis) in the differentiated zone. For induction, seedlings were treated 19 hours with 10 μM of β-estradiol.
