## Supplemental Figure 2 for "Directional Cell-to-cell Transport in Plant Roots"

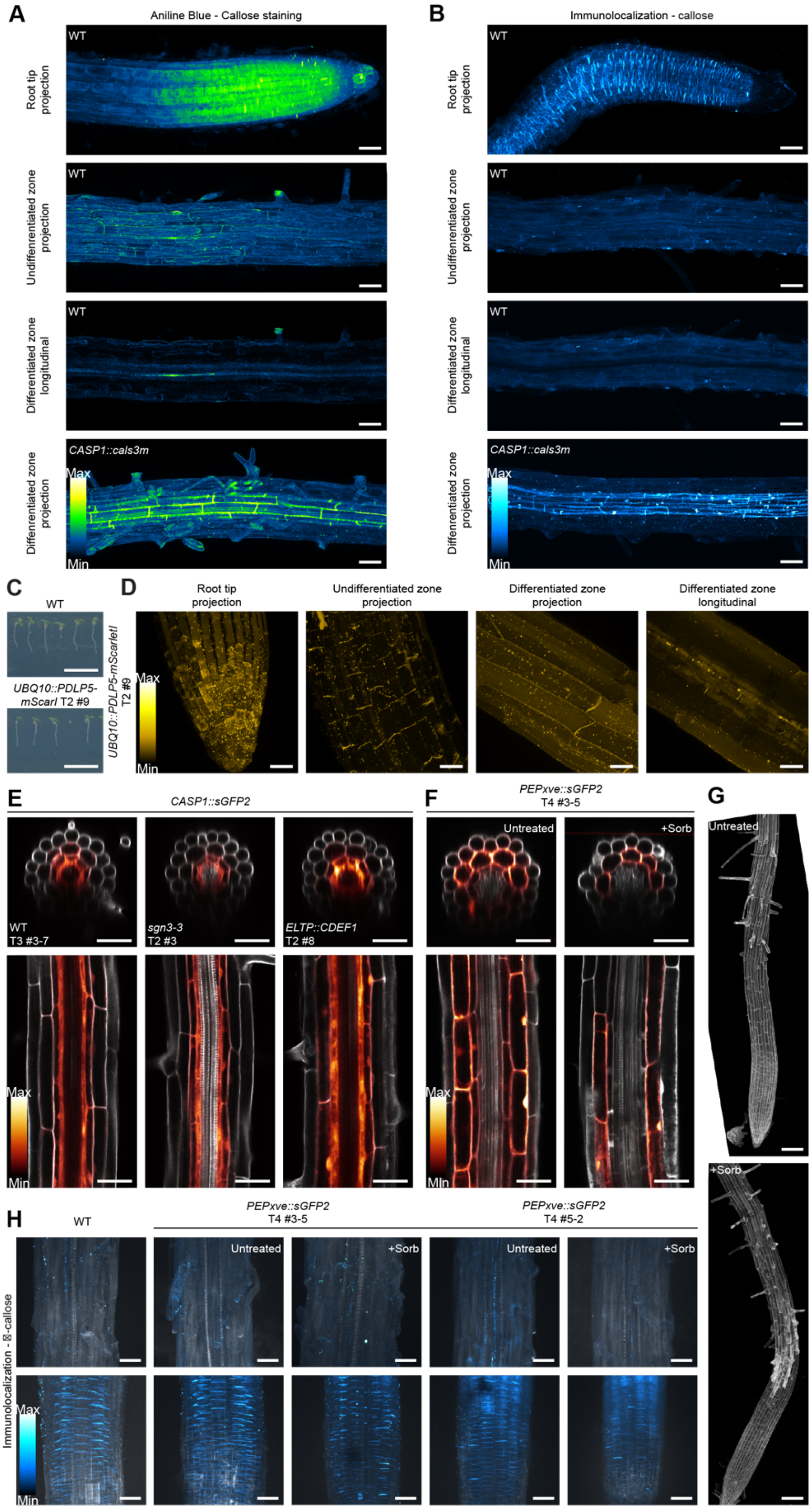


### Figure S2: Water flow affects the symplastic transport in differentiated roots.

(A-B) Callose imaging in WT and *CASP1::cals3m* seedlings stained with aniline blue (A) and after immunolocalization with anti-callose antibody (B) represented as a LUT (Green Fire Blue and Cyan Hot respectively), with 3D projections at the root tip, undifferentiated and differentiated zones or longitudinal views in differentiated zone as indicated in the left. (A, B, E, F) Scale bars, 40 μm. (C) Pictures of 5-day-old seedlings of the WT and *UBQ10::PDLP5-mScarletI*. Scale bars, 10 mm. (D) Live imaging of *PDLP5-mScarletI* expressed under *UBQ10* promoters in root tip, undifferentiated zone and differentiated zone. mScarletI signal is shown as a LUT (Yellow Hot). Scale bars, 20 μm. (E) Live imaging of *CASP1::sGFP2* transgene in WT, *sgn3-3,* and *ELTP::CDEF1* backgrounds in the differentiated zone. (E, F) sGFP2 signal is shown as a LUT (Glow); PI used to stain cell walls is shown in gray, transversal (upper panels) and longitudinal (bottom panels) views are shown. (F) Live imaging of *PEPxve::sGFP2* line under untreated conditions (left panels) and after 2 hours of treatment with 150 mM sorbitol (right panels). For short induction, seedlings were treated with 10 μM of β-estradiol 2 hours prior to the sorbitol treatment. (G) PI staining (tile images with z-projection) of seedlings with similar treatment as in F. Scale bars, 100 μm. (H) Immunolocalization with anti-callose antibody in WT and *PEPxve::sGFP2* lines in root tip (bottom panels) and in differentiated zone (upper panels) for seedlings untreated or after 2 hours of treatment with 150mM sorbitol following 2 hours induction with 10 μM β-estradiol. Immunodetected signal shown as a LUT (Cyan Hot). Scale bars, 30 μm.
