## Supplemental Figure 3 for "Directional Cell-to-cell Transport in Plant Roots"

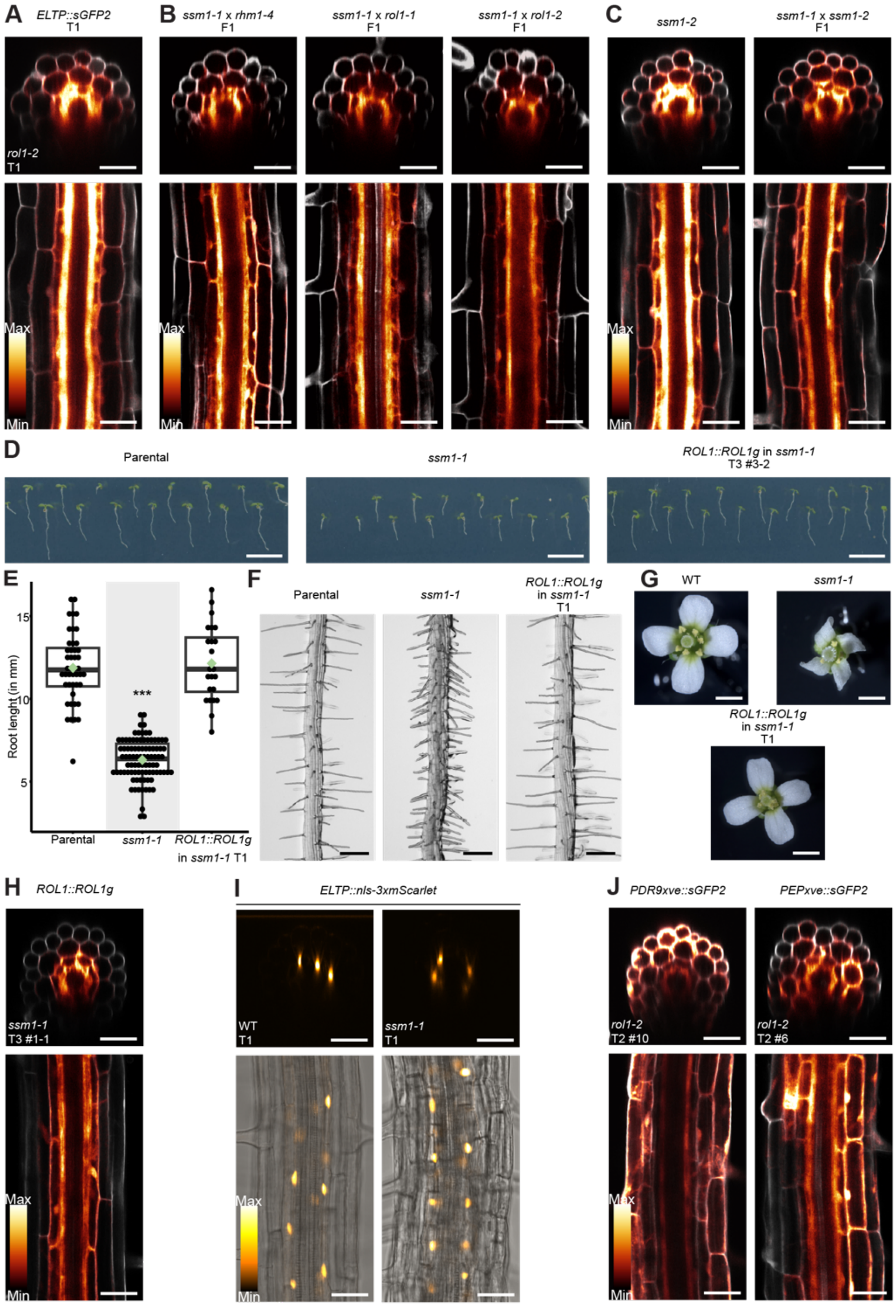


**Figure S3: SESAME genetic screen identified a mutant with bidirectional and exacerbated symplastic transport.**
(A-C) Live imaging of *ELTP::sGFP2* transgene in *rol1-2* (A), F1 seedlings from *ssm1-1* crosses with *rhm1-4*, *rol1-1* and *rol1-2* (B), and in *ssm1-2* and F1 seedlings from crosses between *ssm1-1* and *ssm1-2* (C). (A-C, H, J) Transversal (upper panels) and longitudinal (bottom panels) views are shown; sGFP2 signal is shown as a LUT (Glow); PI used to stain cell walls is shown in gray. Scale bars, 40 μm. (D) Pictures of 5-day-old seedlings of the parental line (*ELTP::sGFP2*), *ssm1-1* and *ROL1::ROL1g* in *ssm1-1*. Scale bars, 10 mm. (E) Quantifications of 5-day-old root length of the parental line (*ELTP::sGFP2*), *ssm1-1* and *ROL1::ROL1g* in *ssm1-1* backgrounds. Data presented as boxplots with dot plots overlaid, representing each individual data point and mean (green diamond), ($n \geq 24$). Differential statistical significance to WT after nonparametric Tukey test are indicated (***, $P < 0.0005$). (F) Pictures of the parental line (*ELTP::sGFP2)*, *ssm1-1* and *ROL1::ROL1g* in *ssm1-1* roots in the differentiated zone. Scale bars, 200 μm. (G) Pictures of WT, *ssm1-1* and *ROL1::ROL1g* in *ssm1-1* flowers from 3-week-old plants. Scale bars, 1 mm. (H) Live imaging of *ELTP::sGFP2* transgene in *ROL1::ROL1g* in *ssm1-1*. (I) Live imaging of *ELTP::nls-3xmScarlet* transgenes in WT (left panels) and *ssm1-1* (right panels) backgrounds in the differentiated zone. Transversal (upper panels) and longitudinal (bottom panels) views are shown. mScarlet signal shown as a LUT (Orange Hot) merged with bright field in longitudinal views. Scale bars, 40 μm. (J) Live imaging of *PDR9xve::sGFP2* (left panels) and *PEPxve::sGFP2* (right panels) transgenes in *rol1-2* background in the differentiated zone. For inductions, seedlings were treated 20 hours with 10 μM β-estradiol.
