## Supplemental Figure 4 for "Directional Cell-to-cell Transport in Plant Roots"

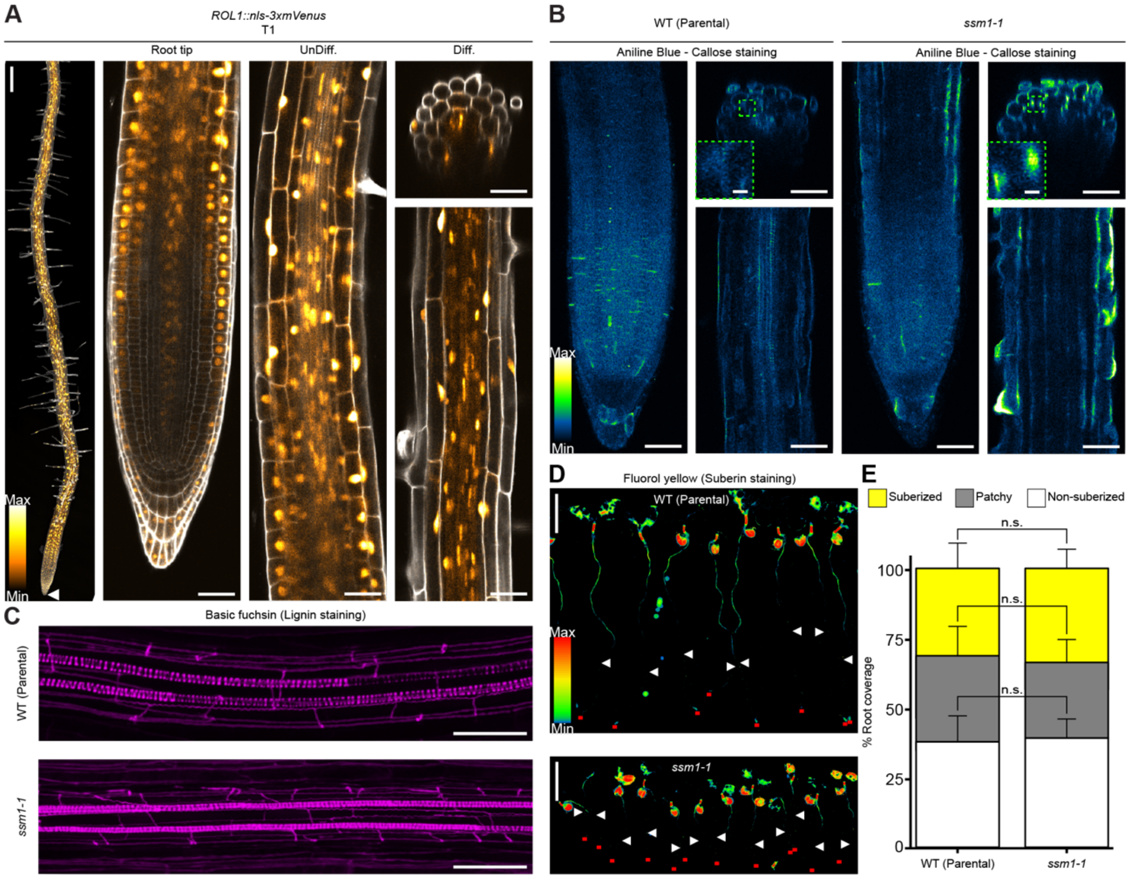


**Figure S4: *ssm1,* a mutant with major cell wall and plasmodesmata defects.**

(A) Live imaging of *ROL1::nls-3xmVenus* promoter-reporter line as a tile image (z-projection, white arrows highlight the signal onset), in the root tip, undifferentiated (UnDiff.) and differentiated (Diff.) zones, mVenus signal is shown as a LUT (Orange Hot); PI used to stain cell walls is shown in gray. Scale bars, 200 µm (tile image) and 40 µm. (B) Callose imaging in WT and *ssm1-1* seedlings stained with aniline blue represented as a LUT (Green Fire Blue), in root tip (left panels) and differentiated zone (transversal view upper right panels and longitudinal view right bottom panel). Scale bars, 40 µm and 4 µm (closer views). (C) Lignin imaging in WT and *ssm1-1* stained with basic fuchsin represented in magenta, in the differentiated zone (z-projection). Scale bars, 40 µm. (D) Suberin imaging in WT and *ssm1-1* seedlings stained with Fluorol yellow represented as a LUT (LSM). The red lines at the bottom mark the root tips and the white arrows the onset of suberization. Scale bars, 2 mm. (E) Quantifications of suberin pattern along the root presented as percentage of the roots corresponding to the undifferentiated, patchy and fully suberized zones. Data represented as overlaid histograms with SD (n ≥ 11). Statistically significant differences between genotypes for each zone of suberization were tested using t.test (n.s. for non-significant).
