## Supplemental Figure 5 for "Directional Cell-to-cell Transport in Plant Roots"

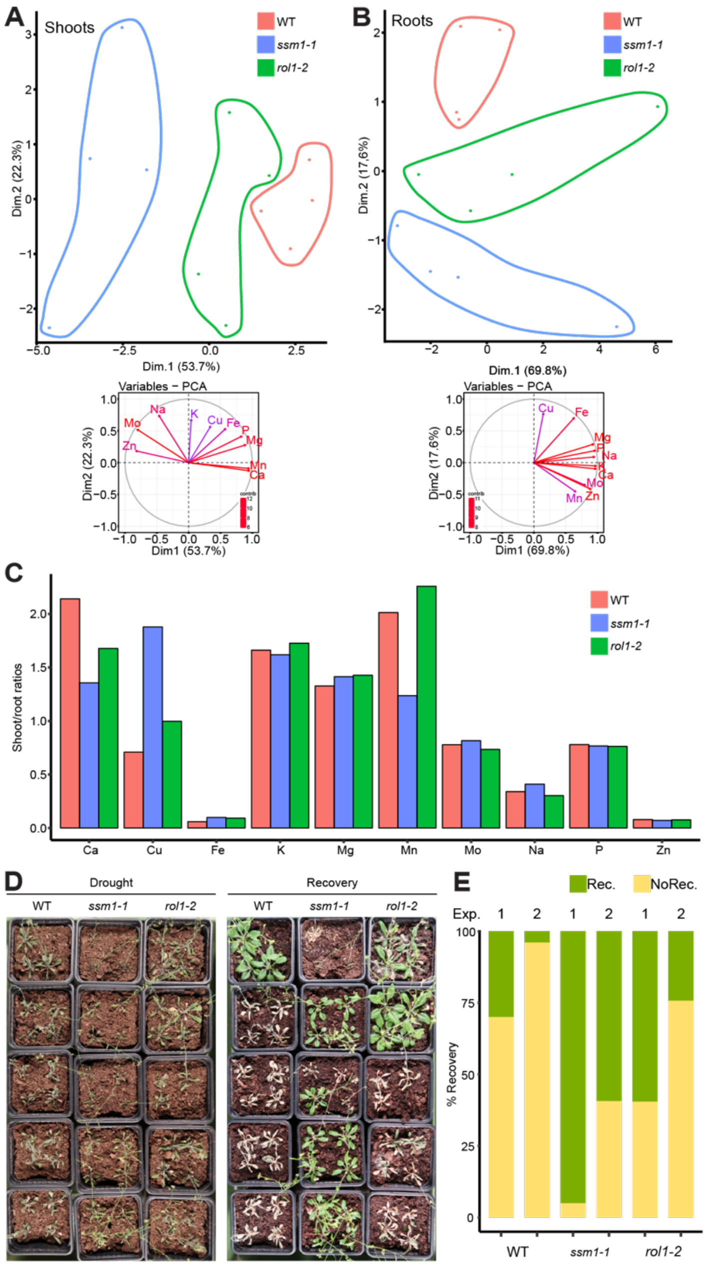


### Figure S5: *ssm1*, a mutant with specific developmental and nutritional defects.

(A-B) PCA plot (upper panel) for the shoots (A) and roots (B) ionomes of WT (in red), *ssm1-1* (in blue) and *rol1-2* (in green) grown 9 days on plates. The x-axis and y-axis represent the two most informative axes (Dim.1 and Dim.2). Variable factor map (bottom panel) representing the amount of variance from each trait on the total variance in the PCA. Longer arrows represent larger amount from total variance. (C) Shoot/root ratio of the different minerals in WT, *ssm1-1* and *rol1-2* to evaluate root to shoot translocation of individual minerals. (See Table S2 for numerical values). (D) Pictures of WT, *ssm1-1* and *rol1-2,* 4-week-old plants grown 2 weeks without watering (Drought condition, left panel) and 4 days after rewatering (Recovery, right panel). (E) Quantifications of the percentage recovery of WT, *ssm1-1* and *rol1-2* plants after drought stress followed by 4 days of rewatering. Two independent experiments (Exp. 1 and 2) are presented (n ≥ 18). Recovered plants (Rec.) are represented in green and non-recovered (NoRec.) plants in yellow.
