## Supplemental Table 1 for "Directional Cell-to-cell Transport in Plant Roots"

**TableS1: Strongest candidate genes list in *ssm1-1* generated by SIMPLE pipeline**

| Chr[1] | position | Ref[2] | Alt[2] | Mutation effect | At_num | CDS change | Protein change |
|---|---|---|---|---|---|---|---|
| 1 | 25340103 | G | A | upstream_gene_variant | AT1G67600 | -2648C>T | |
| 1 | 26225402 | G | A | missense_variant | AT1G69710 | 2254G>A | Val752Met |
| 1 | 27020626 | G | A | missense_variant | AT1G71830 | 853G>A | Gly285Arg |
| 1 | 27280245 | G | A | missense_variant | AT1G72460 | 736G>A | Asp246Asn |
| 1 | 27875520 | G | A | missense_variant | AT1G74130 | 760G>A | Asp254Asn |
| 1 | 28193357 | G | A | splice_region_variant&intron_variant | AT1G75100 | 410+3C>T | |
| 1 | 29299998 | G | A | stop_gained | AT1G77920 | 660G>A | Trp220* |
| 1 | 29550546 | G | A | missense_variant | AT1G78570 | 437G>A | Gly146Asp |

[1]Chromosome, [2]Reference, [3]Alteration
