## Supplemental Table 2 for "Directional Cell-to-cell Transport in Plant Roots"

**TableS2: Raw data for ionomic analysis in shoots and roots (ppm)**

| Genotype | Tissu | Ca | Cu | Fe | K | Mg | Mn | Mo | Na | P | Zn |
|---|---|---|---|---|---|---|---|---|---|---|---|
| WT | Shoots | 2672 | 3.56 | 171.2 | 43626 | 1387 | 101.5 | 1.32 | 1119 | 30265 | 179 |
| WT | Shoots | 2588 | 5.34 | 173.3 | 42129 | 1360 | 100.2 | 1.33 | 1231 | 28711 | 193 |
| WT | Shoots | 2706 | 4.92 | 167.1 | 44082 | 1454 | 102.5 | 1.46 | 1144 | 30872 | 180 |
| WT | Shoots | 2814 | 3.6 | 166.1 | 47599 | 1487 | 103.1 | 1.47 | 1361 | 31288 | 171 |
| *ssm1-1* | Shoots | 1880 | 2.92 | 149.2 | 48028 | 1306 | 82.7 | 2.06 | 1494 | 28213 | 233 |
| *ssm1-1* | Shoots | 1946 | 6.03 | 164.4 | 49781 | 1341 | 83.7 | 2.26 | 1606 | 28478 | 214 |
| *ssm1-1* | Shoots | 1909 | 3.48 | 162.6 | 48916 | 1332 | 81.5 | 1.93 | 1364 | 27893 | 211 |
| *ssm1-1* | Shoots | 1800 | 2.21 | 138.7 | 40734 | 1183 | 77.6 | 2.04 | 1260 | 24832 | 212 |
| *rol1-2* | Shoots | 2321 | 3.1 | 164.7 | 44661 | 1343 | 98 | 1.62 | 1115 | 26723 | 225 |
| *rol1-2* | Shoots | 2182 | 2.77 | 132.2 | 48051 | 1370 | 94.4 | 1.33 | 1047 | 28381 | 192 |
| *rol1-2* | Shoots | 2330 | 3.77 | 177.7 | 52577 | 1389 | 100.3 | 1.53 | 1124 | 29566 | 199 |
| *rol1-2* | Shoots | 2239 | 3.32 | 180.2 | 52316 | 1386 | 98.3 | 1.88 | 1296 | 30660 | 206 |
| WT | Roots | 1186 | 9.49 | 2762.4 | 25727 | 1039 | 44.8 | 2.03 | 3390 | 37843 | 2224 |
| WT | Roots | 1193 | 2.39 | 2959.8 | 25266 | 1030 | 53.6 | 0.96 | 3794 | 38433 | 2389 |
| WT | Roots | 1255 | 3.15 | 2727.8 | 26479 | 1079 | 47.7 | 1.8 | 3316 | 37814 | 2204 |
| WT | Roots | 1400 | 9.53 | 3133.7 | 29275 | 1139 | 56.2 | 2.38 | 3805 | 41384 | 2398 |
| *ssm1-1* | Roots | 1991 | 2.96 | 2179.4 | 41181 | 1302 | 105.6 | 3.95 | 4750 | 49870 | 4478 |
| *ssm1-1* | Roots | 1137 | 1.91 | 1131.8 | 23678 | 798 | 50.2 | 2.54 | 3045 | 31681 | 2719 |
| *ssm1-1* | Roots | 1305 | 1.52 | 1544.9 | 28127 | 860 | 68.4 | 2.28 | 3249 | 33454 | 2820 |
| *ssm1-1* | Roots | 1119 | 1.4 | 1354.8 | 22787 | 691 | 39.1 | 1.4 | 2951 | 27845 | 2323 |
| *rol1-2* | Roots | 1161 | 4.87 | 1304.2 | 24487 | 804 | 41.1 | 1.62 | 3187 | 31250 | 2382 |
| *rol1-2* | Roots | 1270 | 2.41 | 1737.3 | 29596 | 968 | 43.4 | 2.41 | 3745 | 38512 | 2908 |
| *rol1-2* | Roots | 1623 | 2.47 | 2311.3 | 31715 | 1112 | 45.3 | 2.47 | 4456 | 43680 | 2906 |
| *rol1-2* | Roots | 2170 | 4.89 | 3840.8 | 49132 | 1517 | 60.4 | 3.26 | 5811 | 63666 | 4239 |
