## Supplemental Table 3 for "Directional Cell-to-cell Transport in Plant Roots"

**TableS3: Genotyping and cloning primers used in this study, related to STAR Methods**

| Name | Sequence |
|---|---|
| *pEN-L4-PRD9xve-R1* | GATACCGGTGGTACCCAAACGCAGTATAATA |
| | GGAATTGGTACGTACGGCTTGGCTTAAAGA |
| *pEN-L4-PEPxve-R1* | GATACCGGTGGTACCCTCGAGATCAAGACCATCTGTAATCTTTTGA |
| | CCGGAATTGGTACGTACTCGAGGGTTTTGGCTAATGTGATTGTG |
| *pEN-L4-CASP1xve-R1* | GATACCGGTGGTACCCTCGAGTTAATCTGCATAAAAGTGAGTATGAGAGAG |
| | CCGGAATTGGTACGTACTCGAGTTTCTCTTGCAATTGGGGTTTAAAAGA |
| *pEN-L1-3xsGFP2-L2* | CATGGACGAGCTGTACAAGGTGAGCAAGGGCGAGGAG |
| | CTTGTACAGCTCGTCCATGCCGAGAGTG |
| *pEN-L1-mScarlet-L2* | CAACTTTGTACAAAAAAGCAGGCTTAATGGTGAGCAAG |
| | AATGCCAACTTTGTACAAGAAAGCTGGGTTTTACTTGTACAGCTC |
| *pEN-L1-mScarletI-L2* | CAACTTTGTACAAAAAAGCAGGCTTAATGGTGAGCAAG |
| | AATGCCAACTTTGTACAAGAAAGCTGGGTTTTACTTGTACAGCTC |
| *pEN-L1-gPDLP5-L2* | GGCTTACTAGTAAGATCTGCATGATCAAGACAAAGACGACGTCC |
| | TTGAATTCATAAAGGCATGCTTTACACCATTTCTCATCTGCAAAAGGC |
| *pEN-L4-ROL1-R1* | GTTGGATATCAGATCTTCTAGAGGTTTTGAGAGAAATTTCGCGACTCGAG |
| | CTTTTTTGTACAAACTTGCGGATCAAACGCTGCGTTTAAGAAGAAT |
| *pEN-L1-gROL1+600down-L2* | AAAAAGCAGGCTTACTAGTAAATGGCTTCGTACACTCCCAAGA |
| | TTGAATTCATAAAGGCATGCACCTTCAAAGTCGACACATAACAACAAC |
